# Perivascular macrophages collaborate to facilitate chemotherapy resistance in cancer

**DOI:** 10.1101/2022.02.03.478952

**Authors:** Joanne E. Anstee, James W. Opzoomer, Isaac Dean, Henrike P. Muller, Meriem Bahri, Kifayathullah Liakath-Ali, Ziyan Liu, Desmond Choy, Jonathan Caron, Dominika Sosnowska, Richard Beatson, Tamara Muliaditan, Zhengwen An, Cheryl E. Gillett, Guocheng Lan, Xiangang Zou, Fiona M. Watt, Tony Ng, Joy M. Burchell, Shahram Kordasti, David R. Withers, Toby Lawrence, James N. Arnold

**Author notes:** UCL Cancer Institute, University College London, London, WC1E 6DD, United Kingdom. Respiratory Medicine, University College London, London, WC1E 6JF, United Kingdom (Richard Beatson). Department of Pharmaceutical Sciences, Faculty of Science, Utrecht University, Utrecht, Netherlands (Tamara Muliaditan). Central Laboratory of Oral Biomedical Research, Dental Institute, Jilin University, Changchun, China (Zhengwen An). Stem Cell and Regenerative Consortium Centre, LKS Faculty of Medicine, The School of Biomedical Sciences, The University of Hong Kong, Hong Kong, China. Corresponding author: Dr James N. Arnold.

## Abstract

A subset of tumor associated macrophages (TAMs) identified by their expression of the lymphatic vessel endothelial hyaluronan receptor-1 (Lyve-1) reside proximal to blood vasculature and contribute to disease progression. Using a spontaneous murine model of mammary adenocarcinoma (*MMTV-PyMT*), we show that Lyve-1^+^ TAMs, which co-express heme oxygenase-1, form coordinated multi-cellular ‘nest’ structures in the perivascular niche. We show that TAM nest formation is dependent on IL-6 and a communication axis involving CCR5 and its cognate ligands CCL3/4. We demonstrate that Lyve-1^+^ TAM nests are associated with CD8^+^ T-cell exclusion from the tumor and the resistance to immune-stimulating chemotherapeutics. This study highlights an unappreciated collaboration between TAMs and uncovers a spatially driven therapeutic resistance mechanism of these cells in cancer which can be therapeutically targeted.

## Introduction

Macrophages are a phenotypically and functionally diverse population of innate immune cells which become exploited by tumors to facilitate disease progression and therapy resistance (*1*–*8*). Heterogeneity within the tumor associated macrophage (TAM) population arises from their site of origin (*9*) and the influence of environmental cues within the tumor microenvironment (TME) (*3*, *10*, *11*), which can be guided by both spatial (*12*, *13*) and temporal (*11*) parameters. One subset of TAMs reside in close proximity to blood vasculature (<15-20 μm) (*14*–*16*) and are termed perivascular TAMs (PvTAMs). PvTAMs support a variety of pro-tumoral functions including neo-angiogenesis (*15*, *16*), metastasis (*3*, *17*, *18*) and facilitate tumor recurrence post chemotherapy (*19*). Recently, we demonstrated that a subset of PvTAMs expressing lymphatic vessel endothelial hyaluronan receptor 1 (Lyve-1) form a pro-angiogenic Pv niche with a population of pericyte-like cancer associated fibroblasts (CAFs) and orchestrate the platelet-derived growth factor-C (PDGF-C)-dependent expansion of the CAF population with the growing tumor (*16*). Lyve-1 was traditionally considered a marker of lymphatic endothelium (*20*), but has emerged as selective marker for a sub-population of Pv tissue-resident macrophages (*21*–*25*) and TAMs (*16*, *26*). Depleting Lyve-1^+^ PvTAMs in the spontaneous mouse mammary tumor virus polyomavirus middle T antigen (*MMTV-PyMT*) murine model of breast cancer using liposome-based approaches resulted in tumor control, highlighting the importance of the TAM subset in cancer progression and formation of the Pv niche (*16*).

PvTAMs have been demonstrated to develop from a CCR2^+^ monocyte origin (*11*, *27*). These infiltrating monocytes express CXCR4 within the TME in response to tumor-derived tumor growth factor-beta (TGF-β) and subsequently traffic to the Pv space via CXCL12 expressed by a population of PvCAFs (*11*). The CXCR4/CXCL12 axis has also been demonstrated to be important for the accumulation of PvTAMs post chemotherapy treatment (*19*). However, once TAMs reach the endothelium, little is known about the subsequent signaling axes important for niche formation. In the current study we investigate the development of Lyve-1^+^ TAMs and their Pv niche in *MMTV-PyMT* tumors and define a new collaborative function of these cells in forming multi-cellular ‘nests’ which contribute to the resistance to therapeutics through a role in immune exclusion of the TME. This study sheds new light on novel therapeutic strategies to target the immuno-modulatory function of Lyve-1^+^ TAMs.

## Results

### Lyve-1^+^ TAMs express HO-1 and accumulate in Pv ‘nests’ within the TME

Using the autochthonous *MMTV-PyMT* murine model of breast cancer (*28*), we recently demonstrated that Lyve-1 marks a sub-population of pro-angiogenic TAMs in the TME which reside proximal to blood vasculature (*16*) (Fig. 1A). To investigate whether an analogous TAM population exists in human cancer, we extracted the Lyve-1^+^ TAM cell cluster as previously described (*16*) from a scRNA-seq dataset of 9,039 TAMs sorted from 3 individual *MMTV-PyMT* tumors (Fig.1B-C). We mapped the murine Lyve-1^+^ population onto a recently published scRNA-seq atlas for human breast cancer (*29*). The murine Lyve-1^+^ TAM population identified with cells within the human myeloid cell cluster (Fig. 1D and table S1). Focusing on the myeloid cells within the atlas, 1,444 of the 9,675 myeloid cells were judged to be Lyve-1^+^ TAM-like in their phenotype (Fig. 1E-F) and their expression of *Lyve1*, *MRC1* (CD206) and *HMOX1* (HO-1) were significantly associated with the identified cells (Fig. 1F), highlighting a conservation of this TAM phenotype between species. HO-1 is a marker that has previously been associated with PvTAMs in murine models of cancer (*3*, *19*) and the HO-1 expressing TAM population was also observed in the Pv niche of human cancer (Fig. 1G), highlighting a similar spatial location for these cells in human cancer. HO-1 breaks down heme into the biologically active catabolites biliverdin, ferrous iron (Fe^2+^) and carbon monoxide (CO) (*30*, *31*) which has pro-tumoral properties including cytoprotection and immune suppression (*30*, *32*–*37*). HO-1 and Lyve-1 protein co-localized in tumor sections from *MMTV-PyMT* mice (Fig. 1H) and Lyve-1^+^ TAMs were the major source of HO-1 within the Pv niche which also includes, endothelial cells (ECs) and pericyte-like αSMA expressing CAFs (Fig. I-J) (*16*). Interestingly, from these analyses it was evident that the Lyve-1^+^HO-1^+^ PvTAM subset did not uniformly distribute along the endothelium, but instead was restricted to discrete regions of the vasculature where evidence of clustering could be found (Fig. 1A, G, H, J). These clusters, which we herein refer to as ‘nests’, highlighted an unappreciated multi-cellular PvTAM structure within the TME. As Lyve-1 is also expressed at high levels by the lymphatic endothelium (*20*), and HO-1 expression is selectively expressed by this TAM subset (Fig. 1H-J), we generated a knock-in reporter mouse for the *Hmox1* gene to facilitate the study of these cells (Fig. 1K). The reporter consisted of Click Beetle luciferase (Luc) and enhanced green fluorescent protein (eGFP) (*38*) inserted before the stop codon of the genomic *Hmox1* gene locus. HO-1, Luc and eGFP were separated by a self-cleaving P2A sequence (*39*) to allow equimolar expression of the three proteins (Fig. 1K-L and fig. S1A) (mouse herein referred to as HO-1^Luc/eGFP^). As HO-1 plays several important functional roles in the TME (*3*, *31*, *32*, *40*), the arrangement of the reporter construct ensured that the native HO-1 expression was unaffected by the reporter elements. The HO-1^Luc/eGFP^ reporter mouse was crossed onto the *MMTV-PyMT* background and tumors were analyzed for their distribution of HO-1/eGFP expression (fig. S1B). TAMs (fig. S1B), and specifically the Lyve-1^+^CD206^hi^MHCII^lo^ TAM subset (Fig.1M and fig. S2), were the major tumoral source of HO-1/eGFP. Further characterization of F4/80^+^HO-1/eGFP^+^ cells in tumors by immunofluorescence analysis also confirmed the localization of HO-1/eGFP to the Pv space within nest structures (Fig. 1N). These data validate the HO-1^Luc/eGFP^ reporter mouse as a tool for studying Lyve-1^+^ PvTAMs and identifies a previously unappreciated multi-cellular nest structure for these cells in the Pv space.

**Figure 1.**
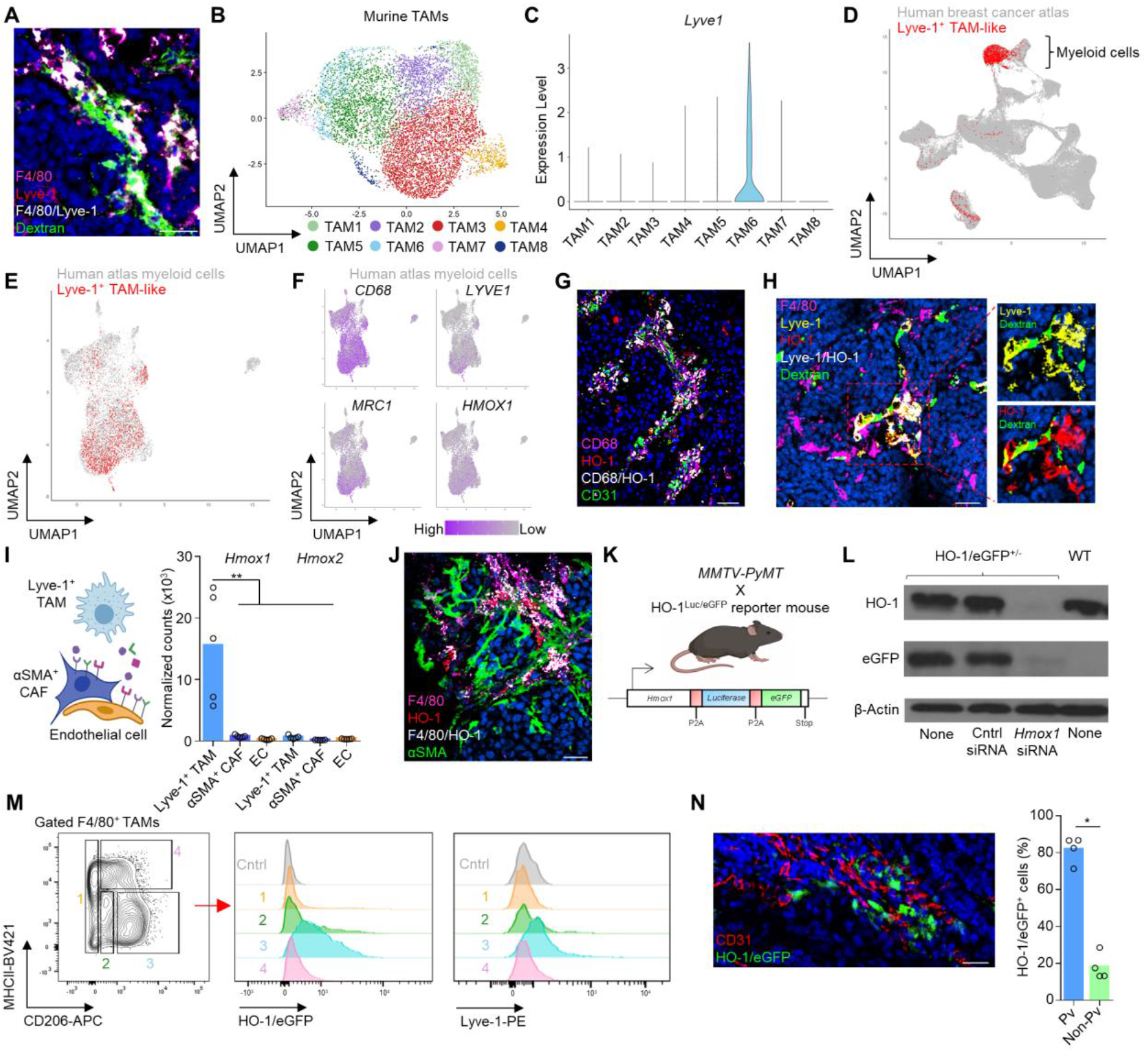
Lyve-1^+^ macrophages co-express HO-1 and reside in nests within the Pv niche. (**A**) Representative image of a frozen section of *MMTV-PyMT* tumor stained with DAPI (nuclei;blue) and antibodies against F4/80 (magenta) and Lyve-1 (red). Functional vasculature was labeled *in vivo* using i.v. dextran-FITC. Colocalizing pixels for F4/80 and Lyve-1 is shown in white. Scale bar is 25 μm. (**B**) UMAP plot of scRNA-seq of F4/80^+^ TAMs from n=3 *MMTV-PyMT* tumors colored by their associated cluster identity, re-analyzed from (*16*). (**C**) Violin plot of Lyve1 expression associated with TAM cluster identity seen in (**B**). (**D**) UMAP plot of scRNA-seq data from a publicly available human breast cancer data set (grey) (*29*) identifying cells (red) that were deemed to display a transcriptional phenotype similar to that of the murine Lyve-1^+^ TAM population (*16*). (**E**) Analysis performed in (**D**) showing specifically the UMAP plot for the myeloid cluster within the atlas. (**F**) UMAP visualizations of *CD68* and selected genes (*LYVE1*, *MRC1*, *HMOX1*) that were preferentially expressed (*P*<2.22×10^-16^) within the Lyve-1^+^-like TAM subset compared to other myeloid cells from the human breast cancer data set (*29*). (**G**) Representative image of a frozen section of human invasive ductal carcinoma stained with DAPI (nuclei;blue) and antibodies against CD68 (magenta), HO-1 (red) and CD31 (green). Colocalizing pixels for F4/80 and HO-1 is shown in white. Scale bar is 50 μm. (**H**) Representative image of a frozen section of *MMTV-PyMT* tumor stained with DAPI (nuclei;blue) and antibodies against F4/80 (magenta), Lyve-1 (yellow) and HO-1 (red). Functional vasculature was labeled *in vivo* using i.v. dextran-FITC. Colocalizing pixels for Lyve-1 and HO-1 is shown in white. Scale bar is 25 μm. (**I**) Schematic depicting the cells in the perivascular niche (left panel) and their relative expression of *Hmox1* and *Hmox2* from bulk RNA-seq (right panel). (**J**) Representative image of a frozen section of *MMTV-PyMT* tumor stained with DAPI (nuclei;blue) and antibodies against F4/80 (magenta), HO-1 (red) and αSMA (green). Colocalizing pixels for F4/80 and HO-1 is shown in white. Scale bar is 25 μm. (**K**) Schematic depicting the transgene of the HO-1-Luc-eGFP knock-in mouse (HO-1^Luc/eGFP^) (lower panel) and the cross that is used in the subsequent tumor studies (upper panel). (**L**) Western blot of HO-1 and eGFP from BMDMs treated with and without *Hmox1* knockdown (KD) siRNA (n=3 repeats). (**M**) Representative contour plot of FACs-gated live (7AAD^-^), CD45^+^F4/80^+^ TAMs from enzyme-dispersed *MMTV-PyMT* HO-1^Luc/eGFP^ tumors separated based on their respective expression of CD206 and MHCII (left panel) and then assessed for their expression of HO-1 and Lyve-1 (right panel; colored histograms) against that of the FMO staining control (grey histogram). Representative of n=10 mice. (**N**) Representative image of a frozen section of a *MMTV-PyMT* HO-1^Luc/eGFP^ tumor stained with antibodies against CD31 (red). Native eGFP fluorescence is shown in green. Scale bar is 25 μm (left panel). Proportion of HO-1/eGFP^+^ cells that are perivascular (<20 μm from live vasculature) across n=4 *MMTV-PyMT* tumors (right panel). Bar charts represent the mean and the dots show individual data points from individual tumors and mice. * *P*<0.05, ** *P*<0.01.

### Lyve-1^+^ macrophages expressing HO-1 can be found populating healthy organs in steady state-conditions

Lyve-1^+^ macrophages reside in a variety of healthy tissues (*16*, *21*–*25*) and *in vivo* bioluminescence imaging of HO-1/Luc in HO-1^Luc/eGFP^ reporter mice demonstrated widespread expression of HO-1 in healthy tissues (Fig. 2A-B). Together this prompted the question whether tissue resident Lyve-1^+^ macrophages share a phenotypic resemblance to the TAM population and co-expressed HO-1. Analysis of healthy tissues from the HO-1^Luc/eGFP^ reporter mouse revealed the spleen, lung, mammary gland, visceral adipose, skin and liver to be some of the highest expressors of HO-1 (Fig. 2B). Flow cytometry analysis of enzyme-digested tissues demonstrated tissue resident F4/80^hi^ macrophages to be the main source of HO-1/eGFP in the healthy tissues analyzed (Fig. 2C). Furthermore, HO-1/eGFP^+^ macrophages expressed Lyve-1 in all organs analyzed apart from the spleen (Fig. 2C-D). However, the spleen represents a unique tissue as it is the major site of erythrophagocytosis (*41*) and heme is a major inducer of HO-1 (*31*). Lyve-1^+^ macrophages in the lung, mammary gland, visceral adipose, skin and liver accounted for almost all HO-1^+^ events in these tissues, highlighting a close concordance between these markers (Fig. 2E). However, despite this similarity, there were distinct differences between the Lyve-1^+^ macrophages between tissues and the tumor in their expression of the markers CD206 and MHCII (Fig. 2D). These data suggest there is a microenvironmental influence on the Lyve-1^+^ macrophage phenotype. Lyve-1^+^ macrophages in healthy tissues arise from a recruited monocyte progenitor (*25*). To confirm the source of Lyve-1^+^ TAMs we utilized the photoconvertible *Kaede* mouse (*42*) crossed to the *MMTV-PyMT* model. When tumors reached 100mm^3^ the TME was photoconverted from *Kaede*-green to *Kaede*-red using a UV-light source (Fig. 2F). After 48 h post photoconversion of the TME there was a clear *Kaede*-green population of Lyve-1^+^TAMs in the TME, indicative of recruitment from a peripheral source, highlighting a monocytic origin for these cells (Fig. 2G). These data highlight a close concordance between Lyve-1 and HO-1 expression associated with this macrophage subset in both healthy tissues and the tumor. Furthermore, we demonstrate that the TME exploits the development of Lyve-1^+^ TAMs through a microenvironmental influence.

**Figure 2.**
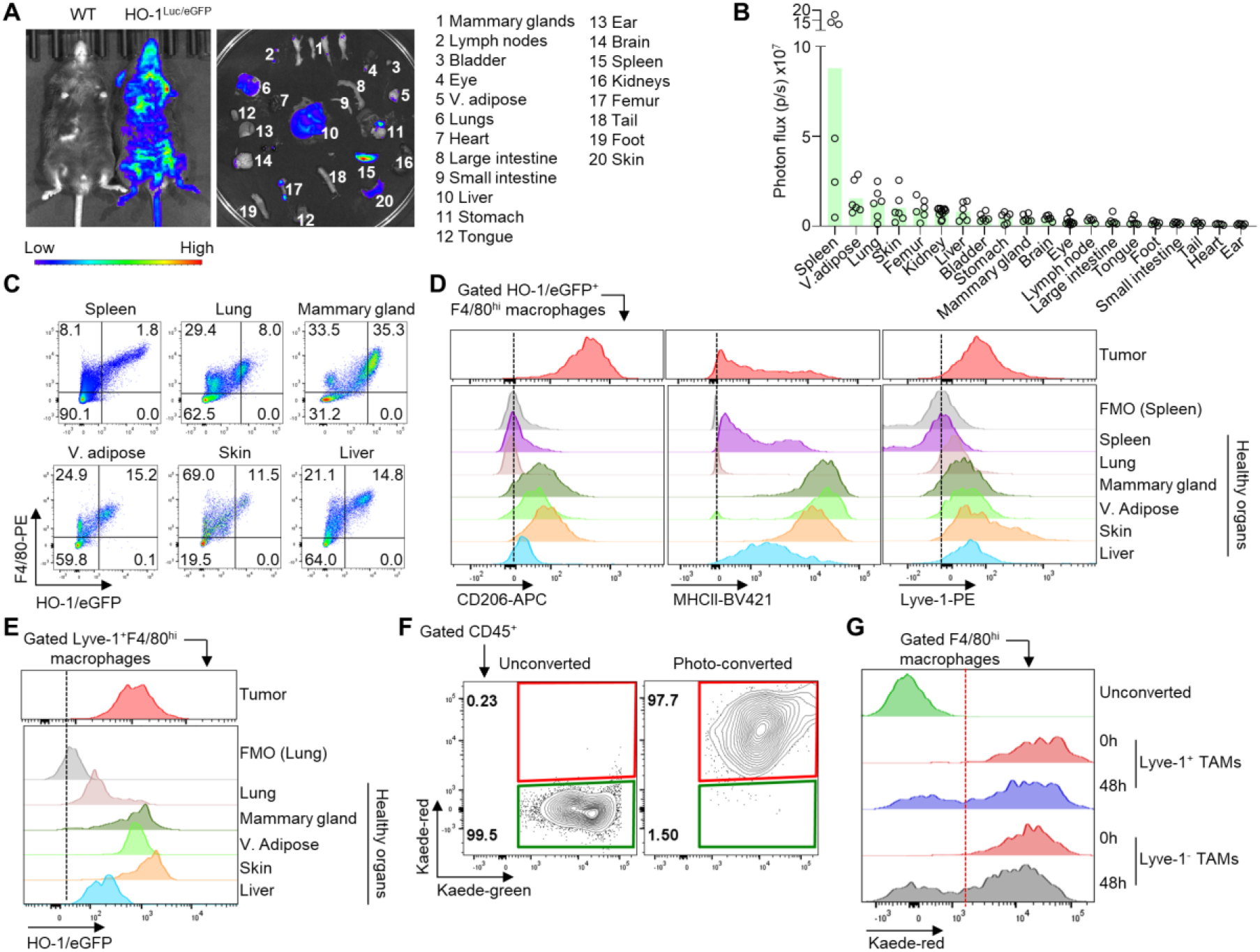
Lyve-1^+^HO-1^+^ macrophages are phenotypically distinct between tissues and the tumor. (**A**) Representative bioluminescence image comparing the relative luciferase expression between a HO-1^Luc/eGFP^ reporter and WT mouse for the whole body (left panel) and dissected indicated tissues (right panel). V.adipose; visceral adipose. (**B**) Quantitated relative Luc expression across the tissues shown in (**A**) from HO-1^Luc/eGFP^ mice (n=6). (**C**) Representative flow cytometry plots of FACs-gated live (7AAD^-^) cells from the indicated tissues of non-tumor bearing HO-1^Luc/eGFP^ mice showing their respective expression of HO-1/eGFP and F4/80. (**D**) Representative histograms of surface CD206 (left panel), MHCII (centre panel) and Lyve-1 (right panel) expression in FACs-gated live (7AAD^-^) F4/80^hi^HO-1/eGFP^+^ macrophages across the indicated healthy tissues shown in (**C**) and an *MMTV-PyMT* HO-1^Luc/eGFP^ tumor for comparison. (**E**) Representative histograms of HO-1/eGFP expression in FACs-gated live (7AAD^-^) F4/80^hi^Lyve-1^+^ macrophages in the indicated healthy tissues and *MMTV-PyMT* tumor. (**F**) Established tumors in *MMTV-PyMT Kaede* mice were photoconverted to *Kaede*-red. (**G**) 48 h after photoconversion tumors were analyzed for *Kaede*-red using flow cytometry, where cells negative for *Kaede*-red represent recruitment from outside the TME. Bar charts represent the mean and the dots show individual data points from individual tumors and mice. * *P*<0.05

### Lyve-1^+^ TAMs are polarized by IL-6 in the TME

To investigate the development of Lyve-1^+^ TAMs within TME, we analyzed scRNA-seq data of TAMs from *MMTV-PyMT*tumors (*16*). Using the QIAGEN Ingenuity Pathway Analysis software (QIAGEN IPA) (*43*), IL-6 signaling was predicted to be an upstream polarization signal associated with the transcriptional programs active in the Lyve-1^+^ TAM subset (Fig. 3A). To investigate the role of IL-6 in polarization of the Lyve-1^+^ PvTAM subset we crossed *MMTV-PyMT* mice onto an *Il6*^-/-^ background and analyzed the tumors in these animals. In the absence of IL-6, tumors were significantly slower to establish, with a median latency of 87 versus 100 days for *Il6*^+/+^ and *Il6*^-/-^ *MMTV-PyMT* mice respectively (Fig. 3B). When tumors reached ~500 mm^3^ they were enzyme-dispersed and analyzed by flow cytometry. Despite no overall change in the abundance of TAMs within the TME (Fig. 3C), there was a significant and selective loss of Lyve-1^+^ TAMs (Fig. 3D-F), highlighting that IL-6 was fundamental to the polarization of the TAM subset. Analysis of bulk RNA-seq data from sorted TME stromal populations within the Pv niche (*16*), highlighted the αSMA pericyte-like CAF and endothelial cells (but not the TAMs) to express *Il6* mRNA (Fig. 3G). More widely, CAFs and endothelial cells were the main tumoral sources of *Il6* mRNA expression with no detectable expression in the tumor cells or other stroma subsets (fig. S3). To identify the spatial location of the *Il6* mRNA in the TME we used RNAscope (Fig. 3H). *Il6* mRNA was detectable in nearly all endothelial cells but high expression of *Il6* mRNA was evident in those endothelial cells that were proximal to HO-1^+^ cells (Lyve-1^+^ TAMs) (Fig. 3H and fig. S4), although αSMA^+^ CAFs could be found expressing *Il6* mRNA in the TME, as indicated by the bulk RNA-seq data, the cells expressing *Il6* mRNA were not proximal to the HO-1^+^ cells (fig. S4). These data suggest that Lyve-1^+^ TAMs develop in response to IL-6 that is secreted by the endothelium in the TME.

**Figure 3.**
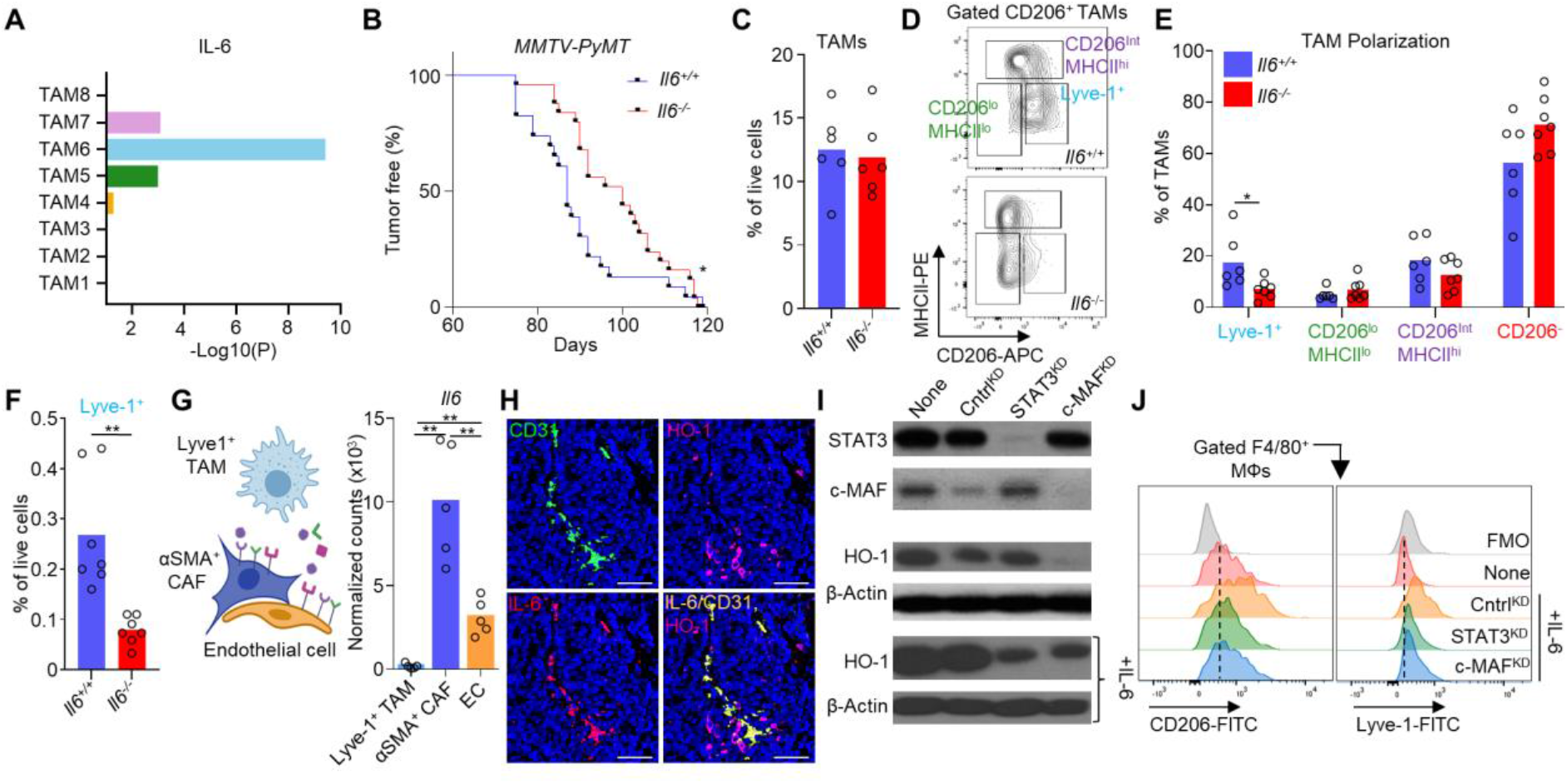
Lyve-1^+^ PvTAMs are polarized by IL-6 in the tumor. (**A**) Plot showing the significance value of IL-6 as an enriched upstream regulator across the TAM populations identified in Fig. 1B. (**B**) Kaplan-Meier curve showing the tumor latency period for *Il6*^+/+^ (WT; n=23) and *Il6*^-/-^ (n=25) *MMTV-PyMT* mice. (**C**) Abundance of live (7AAD^-^) CD45^+^F4/80^+^Ly6C^-^ TAMs in enzyme-dispersed tumors from *Il6*^+/+^ and *Il6*^-/-^ *MMTV-PyMT* mice assessed using flow cytometry (n=6 per group). (**D-F**) Representative contour plot of FACs-gated live (7AAD^-^), F4/80^+^CD206^+^ TAMs from enzyme-dispersed *MMTV-PyMT* tumors showing their respective expression of CD206 and MHCII in *Il6*^+/+^ (upper panel) and *Il6*^-/-^ (lower panel) mice (**D**) and quantification of the respective TAM populations (**E**) and abundance of Lyve-1^+^ TAMs as % of live cells (**F**) (n=7 per group). (**G**) Schematic depicting the cells in the Pv niche (left panel) and their relative expression of *Il6* mRNA from bulk RNA-seq (right panel). (**H**) Representative image of a FFPE section of *MMTV-PyMT* tumor stained with DAPI (nuclei;blue) and antibodies again CD31 (green), HO-1 (magenta) and probed for *ll6* mRNA (red). Scale bar is 25 μm. (**I**) Western blots of STAT3 and c-MAF from BMDMs treated with and without IL-6 and the indicated knockdown (KD) siRNAs (n=3 repeats). (**J**) Representative histograms of surface CD206 (left panel) and Lyve-1 (right panel) expression assessed using flow cytometry of BMDMs treated as described in (**I**) (n=3 repeats). Image in panel (**G**) was created using BioRender software. Bar charts represent the mean and the dots show individual data points from individual tumors and mice. * P<0.05, ** P<0.01.

### IL-6 polarizes Lyve-1^+^ macrophages via a STAT3/c-MAF-dependent signal

To investigate whether IL-6 stimulation alone was sufficient to generate a Lyve-1^+^ TAM-like phenotype. Bone marrow-derived macrophages (BMDMs) were exposed to IL-6 and their phenotype was assessed. IL-6 stimulated BMDMs upregulated the expression of the three key phenotypic markers; HO-1 (Fig. 3I and fig. S5), CD206 and Lyve-1 (Fig. 3J) which define the subset. These data suggested that IL-6 plays a dominant role in the polarization identity of Lyve-1^+^ TAMs and their relative absence in *Il6*^-/-^ *MMTV-PyMT* mice (Fig. 3F) was most likely directly due to the loss of IL-6 from the TME. Recent data demonstrated that the transcription factor c-MAF was vital to the development of Lyve-1^+^ Pv macrophages in healthy tissues (*44*), and we sought to establish whether c-MAF signaling may account for the Lyve-1^+^ macrophage phenotype observed through IL-6 stimulation. IL-6 signaling is associated with the JAK/STAT3 pathway (*45*), however, STAT3 has been linked to the expression of c-MAF via the basic leucine zipper transcription factor ATF-like, Batf in T follicular helper cells (*46*). To investigate whether c-MAF and STAT3 signaling were required for the Lyve-1^+^ macrophage phenotype, we knocked down the expression of either c-MAF or STAT3 in BMDMs prior to stimulation by IL-6 (Fig. 3I and fig. S5). Loss of either transcription factor was sufficient to prevent the upregulation of Lyve-1, CD206 and HO-1 upon IL-6 stimulation (Fig. 3I and J and fig. S5). Interestingly, c-MAF appeared to be required for even basal HO-1 expression in BMDMs (Fig. 3I and Fig. S5B). These data demonstrate that IL-6 is sufficient to generate the key phenotypic markers associated with the Lyve-1^+^ PvTAM subset through a STAT3 and c-MAF-dependent pathway.

### Lyve-1^+^ TAMs communicate via a CCR5-dependent axis to orchestrate Pv nest formation

Having identified IL-6 as a driver for the Lyve-1^+^ TAM polarization program (Fig. 3), we investigated whether their polarization program might influence a communication axis between these cells to orchestrate the formation of nests within the Pv niche (Fig.1A). A microarray analysis was performed on monocyte-derived macrophages exposed to IL-6, IL-4 (M2) or IFNγ/LPS (M1) (fig. S6A-B). M1 and M2 conditions were included as comparator groups for the polarization extremes of these cells (*47*). The IL-6 polarization program was a distinct program from that of M1/M2 macrophages (fig. S6A-B). Interestingly, IL-6 polarized macrophages (M_(IL-6)_) were associated with higher *Pdgfc* expression (fig. S6C), a growth factor we recently demonstrated to play a role in the Lyve-1^+^TAM-dependent expansion of αSMA CAFs within the Pv niche (*16*). In probing for IL-6 induced chemokine receptors we identified a unique upregulation of the chemokine receptor gene *Ccr5* (fig. S6D). Analyzing the bulk RNAseq data for cells in the Pv niche revealed both *Ccr5* (Fig. 4A) and its cognate ligands *Ccl3* and *Ccl4* (Fig. 4B) to be expressed by the Lyve-1^+^ TAMs within the Pv niche and provided a specific communication axis for these cells. The expression of CCR5 was demonstrated at the protein level in Lyve-1^+^ TAMs from *MMTV-PyMT* tumors (Fig. 4C) and IL-6 stimulated BMDMs (Fig. 4D). Although all TAMs expressed some CCR5, Lyve-1^+^ TAMs were the highest expressor in the TME (fig. S6E). CCR5 was functional on the macrophages and IL-6 stimulated BMDMs could migrate towards the CCR5 ligand CCL5 (Fig. 4E). When IL-6 stimulated BMDMs were also placed in close contact *in vitro* their ability to spread from one another was also reduced (Fig. 4F). To test the role of CCR5 in maintaining Lyve-1^+^ TAMs in nests within the TME we crossed *MMTV-PyMT* mice onto a *Ccr5*^-/-^ background. In the absence of CCR5, tumors were significantly slower to establish, with a median latency of 87 versus 102 days for *Ccr5*^+/+^ and *Ccr5*^-/-^ *MMTV-PyMT* mice respectively (Fig. 4G). As CCR5 expression was downstream of IL-6 signaling in the Lyve-1^+^ TAM subset, as predicted, the absence of CCR5 did not affect the total prevalence of TAMs (Fig. 4H) or the Lyve-1^+^ TAM subset (Fig. 4I-J) in these tumors. However, immunofluorescence imaging of tissue sections from these tumors revealed an increase in the median distance of the Lyve-1^+^ TAMs to the endothelium and each other (Fig. 4K), without effecting their abundance overall (Fig. 4L). This demonstrated that CCR5 was required to maintain the nest structures of Lyve-1^+^ TAMs. To demonstrate that CCR5 also played an ongoing active role in maintaining the Lyve-1^+^ TAM nests post formation, rather than just the initial formation of the structure, we injected *MMTV-PyMT* mice with maraviroc (Fig. 4M), a drug that is clinically used to inhibit CCR5 (*48*). Therapeutically blocking CCR5 signaling using maraviroc in tumors with formed Lyve-1^+^ PvTAM nests did not affect tumor growth (fig. S7) but did result in an observable dispersion of the Lyve-1^+^ TAM nests away from the Pv space (Fig. 4N), highlighting that CCR5 represents an ongoing communication axis for the maintenance of the Pv nest structures. These data highlight a new role for CCR5 in the collaborative formation and maintenance of the Lyve-1^+^ TAM nests in the Pv niche.

**Figure 4.**
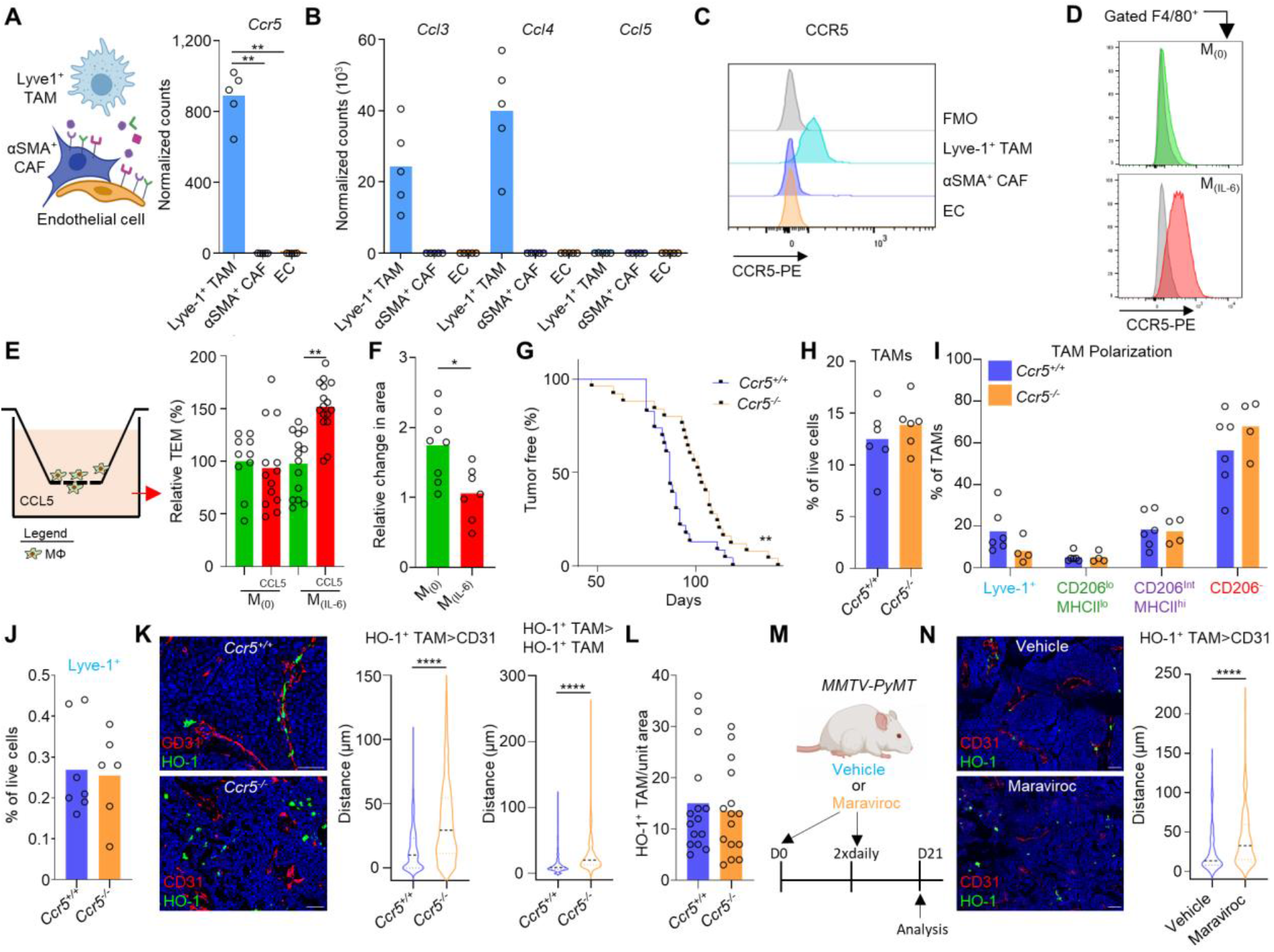
Lyve-1^+^ TAMs communicate via a CCR5 dependent axis to orchestrate Pv nest formation. (**A**) Schematic overview of cells in the Pv nest (left panel). Bar plot depicting normalized gene expression values for *Ccr5* in the bulk RNA-seq of the Pv niche populations across n=5 mice. (**B**) Bar plot depicting normalized gene expression values for *Ccl3*, *Ccl4* and *Ccl5* in the bulk-RNA-seq described in (**A**) across n=5 mice. (**C**) Representative histograms of surface CCR5 expression for the indicated populations, against that of the fluorescence minus one (FMO) staining control (grey histogram) in enzyme-dispersed *MMTV-PyMT* tumors using flow cytometry. αSMA^+^ CAFs are gated based on CD45^-^CD90^+^CD34^-^ as described (*16*). (**D**) Representative histograms of surface CCR5 expression in BMDMs treated with (red filled histogram lower panel) and without (green filled histogram upper panel) IL-6, against that of the FMO staining (grey filled histogram) (n=5 repeats). (**E**) Schematic of the assay (left panel). Relative transendothelial migration (TEM) of M_(0)_ and M_(IL-6)_ BMDMs in the presence or absence of CCL5 (right panel) (n=10-16 where each dot represents an individual insert). (**F**) Relative change in the area of plated M_(0)_ and M_(IL-6)_ BMDMs spheroids over a 72 h. Each point represents an individual spheroid. (**G**) Kaplan-Meier curve showing the tumor latency for *Ccr5*^+/+^ (WT; n=23) and *Ccr5*^-/-^ (n=25) *MMTV-PyMT* mice. (**H-J**) Tumors from *Ccr5*^+/+^ and *Ccr5*^-/-^ *MMTV-PyMT* mice were enzyme-dispersed and assessed using flow cytometry for the abundance of live (7AAD^-^) CD45^+^F4/80^+^Ly6C^-^ TAMs (**H**), and their phenotype as proportions of the TAM gate (**I**) and the abundance of Lyve-1^+^ TAMs as a % of live cells (**J**) (n=6-7 per group). (**K**) Representative images of a frozen section of *MMTV-PyMT* tumor stained with DAPI (nuclei;blue) and antibodies against CD31 (red) and HO-1 (green) in *Ccr5*^+/+^ (upper panel) and *Ccr5^-/^-* (lower panel) *MMTV-PyMT* mice. Scale bar is 50 μm (left panel). Distance of individual HO-1^+^ TAMs to the nearest CD31^+^ cell (middle panel) and HO-1^+^ TAMs to the nearest HO-1^+^ TAM (right panel) in *Ccr5*^+/+^ and *Ccr5^-/-^ MMTV-PyMT* tumors (taken from multiple images from n=5 tumors per group). (**L**) Relative abundance of HO-1^+^ TAMs per unit area as assessed by immunofluorescence imaging in *Ccr5*^+/+^ and *Ccr5*^-/-^ *MMTV-PyMT* tumors (taken from multiple images from n=5 tumors per group). (**M**) Schematic representing the dosing strategy for the CCR5 inhibitor maraviroc and vehicle in *MMTV-PyMT* mice (n=3 per group). (**N**) Representative images of a frozen section of *MMTV-PyMT* tumor stained with DAPI (nuclei;blue) and antibodies against CD31 (red) and HO-1 (green) in vehicle treated mice (upper panel) and maraviroc treated mice (lower panel). Scale bar is 50 μm (left panel). Distance of individual HO-1^+^ cells to nearest CD31 staining across vehicle and maraviroc treated *MMTV-PyMT* tumors (taken from multiple images from n=3 tumors per group). Image in panel (**A**) was created using *BioRender* software. Bar charts represent the mean and the dots show individual data points from individual tumors and mice. * *P*<0.05, ** *P*<0.01, \*\*\*\**P*<0.0001.

### Lyve-1^+^ TAM nests support immune exclusion in the TME

To understand how the absence of Lyve-1^+^ TAM nests may alter the wider TME we analyzed the composition of the stroma in ~500 mm^3^ tumors from WT, *Il6*^-/-^ and *Ccr5*^-/-^ *MMTV-PyMT* mice (Fig. 5A). The stromal composition in enzyme-dispersed tumors were assessed using flow cytometry (Fig. 5B and fig. S8). Broadly, the stromal composition of the tumors was highly similar despite the spontaneous nature of the tumor model (Fig. 5B). However, the only consistent difference for both the *Il6*^-/-^ and *Ccr5*^-/-^ *MMTV-PyMT* mice compared to WT tumors was a significant increase in tumor infiltrating CD8^+^ T-cells (Fig. 5B). The CD8^+^ T-cells had a similar overall proportion of cells which displayed effector function (Fig. 5C and fig. S9). This suggested that the Lyve-1^+^ TAM nests could be associated with immune exclusion in the TME. The endothelium expresses adhesion molecules which permit leukocyte rolling, migration and arrest prior to diapedesis into inflamed tissues (*49*), and it was possible that Lyve-1^+^ TAMs could modulate the endothelium due to their close proximity to one another. However, there was no evidence for a change in the expression of endothelial VCAM-1, ICAM-1 or pNAD in the absence of Lyve-1^+^ TAMs or their nests in the TME (fig. S10A). To explore if Lyve-1^+^ TAMs could play a role in immune exclusion, we established an *in vitro* assay for creating artificial Pv nests (Fig. 5D). In this assay, M_(IL-6)_ which are analogous to the Lyve-1^+^ TAMs (Fig.3I-J), were seeded onto the basolateral side of a transwell insert, and then a basement membrane and an endothelial layer were seeded on the apical side of the insert (Fig. 5D). The presence of M_(IL6)_ had no significant effect on the permeability of the endothelial layer (Fig. 5E). However, when CD8^+^ T-cells were placed in contact with the endothelial layer with a gradient of the T-cell chemokine CXCL10, there was a significant reduction of T-cell migration across the endothelial layer in the presence of M_(IL6)_ (Fig. 5F). HO-1 has been demonstrated to play a role in vascular biology (*31*) and, as such, we considered whether the enzyme might play a direct role in the mechanism of CD8^+^ T-cell exclusion. Pharmacologically inhibiting HO-1 activity using the inhibitor tin mesoporphyrin (SnMP) (*50*) (Fig. 5G), or a genetic knock out in M_(IL6)_ using BM from a *Hmox1*^fl/fl^ mouse crossed with *Lyz2* promoter driven Cre recombinase (Fig. 5H and fig. S10B) resulted in a restoration of T-cell transendothelial migration in the *in vitro* assay (Fig. 5G-H). These data suggest that HO-1 activity could play a role in T-cell exclusion. These *in vitro* data suggest that Lyve-1^+^ TAMs may serve as gatekeepers in dictating CD8^+^ T-cell entry into the tumor (Fig. 5B).

**Figure 5.**
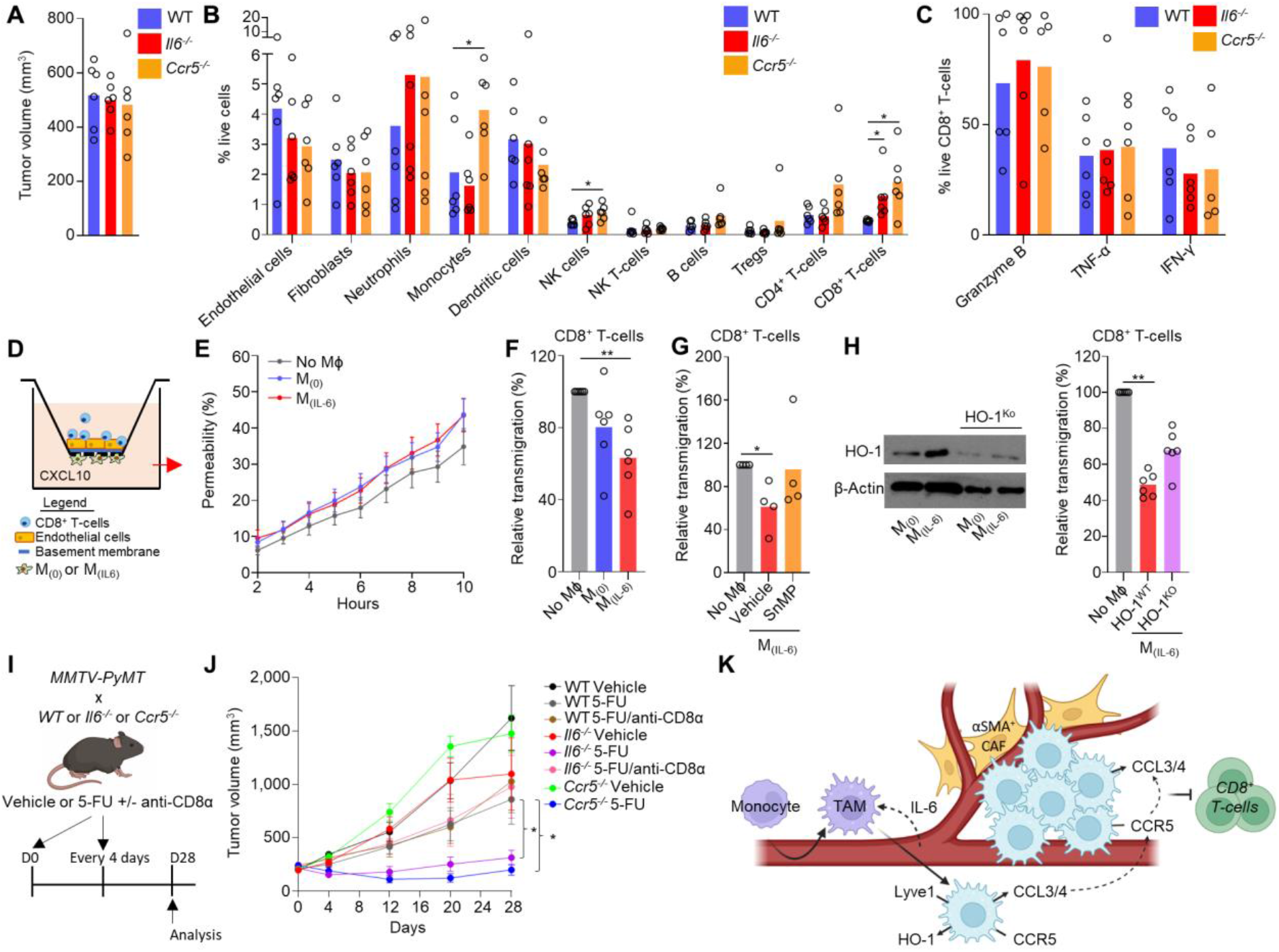
Lyve-1^+^ TAMs and their nests support immune exclusion and resistance to chemotherapy. (**A-C**) Tumors from WT, *Il6*^-/-^ or *Ccr5*^-/-^ *MMTV-PyMT* mice were analyzed for their stromal composition using flow cytometry analysis. Tumor sizes on day of analysis (**A**) and the relative proportions of live (7AAD^-^) stromal cell populations (**B**), are shown. The markers used to differentiate cells can be found in the Materials and Methods section. Gated CD8^+^CD3^+^ T-cells were assessed for their expression of the indicated effector molecules post PMA/ionomycin treatment (**C**). (**D-H**) An *in vitro* Pv niche assay was established to investigate the role of Pv macrophages in CD8^+^ T-cell transendothelial migration. (**D**) A schematic of the assay. Permeability of the endothelial cell layer at the indicated time points to albumin in the presence or absence of M_(0)_ or M_(IL-6)_ cells (**E**). The relative transendothelial migration of CD8^+^ T-cells in the presence or absence of M_(0)_ or M_(IL-6)_ cells on the basolateral surface (**F**) and the effect of the addition of an HO inhibitor SnMP (**G**) and genetic knockout of HO-1 in M_(IL-6)_ (**H**), where the western (left) shows the relative detectable expression of HO-1 in WT and HO-1^KO^ macrophages (using *Hmox1*^fl/fl^ × *Lyz2*^cre^ mice). (**I**) Schematic representing the dosing strategy for 5-FU and/or immune-depleting anti-CD8α antibodies in *MMTV-PyMT* mice. (**J**) Growth curves of established spontaneous tumors in WT, *Il6*^-/-^ and *Ccr5*^-/-^ *MMTV-PyMT* mice that were given 5-FU (40 mg/kg/4 days) or their respective vehicles and immune-depleting anti-CD8α IgG where indicated. Indicated dosing started at day zero (cohorts of n=6-9 mice). (**K**) Schematic overview of the Pv niche. Image created using Biorender. Bar charts show the mean and the dots show individual data points from individual tumors and mice. Line charts display the mean and SEM. * *P*<0.05, ** *P*<0.01.

### Targeting Lyve-1^+^ TAM nests can restore the sensitivity of chemotherapeutics

There is a clear clinical link between the presence of tumor infiltrating lymphocytes (TILs) and the response to therapeutics which can modulate their effects through the anti-tumor immune response, such as immune checkpoint blockade (*51*, *52*) and certain cytotoxic chemotherapies (*53*, *54*). We have demonstrated HO-1 activity to be a major immune suppressive pathway preventing chemotherapy-elicited immune responses in the TME (*55*). We considered whether the presence of Lyve-1^+^ TAMs (the major tumoral source of HO-1), or their nest structures, might be implicated in the resistance to immune modulating cytotoxic chemotherapeutics. Treatment of tumor bearing *MMTV-PyMT* mice with 5-fluorouracil (5-FU), a chemotherapeutic which has been used in the clinic (56) and which has been demonstrated to have immune-stimulatory capabilities (*57*, *58*), did not significantly affect tumor growth in WT mice (Fig. 5I-J). However, when 5-FU was administered to either *Il6*^-/-^ or *Ccr5*^-/-^ mice a significant control of tumor growth was achieved (Fig. 5J). To investigate whether the tumor control was immune-dependent, we depleted of CD8^+^ T-cells using immune-depleting antibodies *in vivo* prior to the initiation of 5-FU treatment in *Il6*^-/-^ *MMTV-PyMT* mice in which the Lyve-1^+^ TAM subset was unable to polarize (fig. S11). In the absence of CD8^+^ T-cells, 5-FU had no effect on tumor growth in *Il6*^-/-^ tumors demonstrating an immunological basis for the mechanism of tumor control observed. As such, these data highlight a previously unappreciated link between Lyve-1^+^ PvTAM nests and the immune landscape of the TME which can provide a resistance mechanism for the immune-mediated effects of cytotoxic chemotherapeutics.

## Discussion

In this study we describe a coordinated and collaborative role for Lyve-1^+^ PvTAMs in forming multi-cellular nest structures within the TME which facilitate immune exclusion and the resistance to the immune-stimulating capabilities of cytotoxic chemotherapy. Although a number of pro-tumoral processes have been described to PvTAMs (*15*), this study highlights an unappreciated mechanism of these cells that is reliant on their collaborative interaction and collusion which promotes cancer progression. This study sheds new light on the development of the Lyve-1^+^ TAM subset. Using photolabeling approaches we demonstrate that Lyve-1^+^ TAMs develop from recruitment of a progenitor into the TME, consistent with previous studies that have demonstrated PvTAMs to have derived from a monocyte origin (*11*, *25*, *27*). It is apparent that PvTAMs develop through a sequential signaling program involving their upregulation of CXCR4 in response to tumor cell-derived TGF-β which guides their migration back to the endothelium on a gradient of CXCL12 expressed by a PvCAF population (*11*). The importance of the CXCR4/CXCL12 axis in PvTAM accumulation at the vasculature has been demonstrated in several studies (*19*, *59*). We propose that the mechanisms presented in this study provide insight on the subsequent developmental step of PvTAMs into the Lyve-1^+^ TAM population post reaching the endothelium. This is supported by the observation that loss of IL-6 resulted in a striking and specific loss of the Lyve-1^+^ TAM population leaving other TAM polarization states unchanged, suggesting it is important for only the terminal step in polarization. Interestingly, the interaction of angiopoietin-2 expressed on endothelial cells and Tie2 expressed on PvTAMs has also been demonstrated to play an important role in their accumulation at the vasculature (*60*, *61*), however we did not find evidence of *Tie2/Tek* gene expression in the RNA-seq datasets for the Lyve-1^+^ TAM in this study.

We demonstrate that a key feature of the Lyve-1^+^ TAM program was high expression of the enzyme HO-1, a gene which we have previously linked to IL-6 signaling (*3*). However, surprisingly, in homeostatic tissues, tissue resident Lyve-1^+^ macrophage populations also expressed HO-1. We identified *in vitro* that c-MAF signaling was required for Lyve-1, HO-1 and CD206 expression on BMDMs in response to IL-6 and could link these markers on tissue resident macrophages in non-inflammed healthy tissues which have recently been demonstrated also to be c-MAF-dependent (*44*). Although the signal for c-MAF in healthy tissues is unknown, it could explain the unexpected high expression of HO-1 which is generally considered as a stress- or inflammation-inducible enzyme (*30*, *62*). In this study, using HO-1^Luc/eGFP^ reporter mice, we find that the Lyve-1^+^ tissue resident macrophages account for almost all HO-1 expression in healthy tissues which highlights a role for macrophages in the homeostatic functions ascribed to HO-1 in healthy tissues (*31*).

The identification of the immunological resistance mechanisms to cytotoxic chemotherapy responses is important as it has become apparent that the immune-stimulating properties of these drugs may underlie a significant proportion of their anti-tumoral efficacy (*63*–*68*). This study also complements the wider association between TAMs representing a pivotal role in resistance to chemotherapy (*6*–*8*, *55*) and facilitating relapse after the cessation of treatment (*19*). This study also helps mechanistically supporting the link between IL-6 suppression and the immune-stimulating effects of cytotoxic chemotherapies (*69*). There are many studies which have described HO-1 as having pro-tumoral properties having important roles in cytoprotection and immune suppression (*30*, *32*–*37*). We previously demonstrated that HO-1 plays a major role in the suppression of anti-tumor CD8^+^ T-cell responses elicited by chemotherapy in *MMTV-PyMT* mice (*55*). Many of the effects of HO-1 have been attributed to its production of CO as a catabolite of heme degradation which can modulate several signaling pathways including p38 MAPK (*70*), STAT1/3 (*71*) and NFκB (*72*, *73*). As such, HO-1 activity can compromise anti-tumor CD8^+^ T-cell responses in the TME (*32*). The superior tumor control observed when Lyve-1^+^ TAMs, which are the exclusive tumoral source of HO-1, could not develop their nest structures (*Ccr5*^-/-^), highlight a spatial parameter associated with its immune-suppressive capabilities and further emphasises the collaborative nature of this suppression. It has been demonstrated that macrophage density can also relate to a ‘quorum licensing’ of macrophage activation (*74*), it would be interesting to understand how the nest structures might also influence or refine the effector function of Lyve-1^+^ TAMs in the context of cancer.

It is clear that PvTAMs reside in unique niche arrangements to support their function, such as the described TMEs of metastasis” (TMEM), where a PvTAM, tumor cell expressing a slice variant of mammalian-enabled protein ‘Mena’ (*75*), and an endothelial cell are in direct contact to facilitate transendothelial migration of tumor cells into the blood from the tumor (*17*, *76*–*79*). However, the heterogeneity of PvTAMs still requires further investigation, as not all PvTAMs expressed Lyve-1 which may represent a progenitor stage or discrete subset of these cells. In this study we characterize the functionality associated with a multi-cellular Lyve-1^+^ PvTAM structure and define a new collaborative action of these cells to form multi-cellular biological units which are associated with immune exclusion of CD8^+^ T-cells from the TME. Interestingly, PvTAMs have been demonstrated to play an active role in neutrophil recruitment to inflamed skin in response to *Staphylococcus aureus* infection (*80*), suggesting that PvTAMs may play a gatekeeper role to modulate the immune landscape of the TME, however in *MMTV-PyMT* tumors the loss of Lyve-1^+^ TAMs, or their nests, resulted in a specific increase in the abundance of only CD8^+^ T-cells in the TME. An IL-6 driven CCR5 expression by Lyve-1^+^ TAMs provide the means to connect a TAM>TAM communication axis of CCR5-CCL3/4 which maintained their nest structures. Interestingly, blockade of CCR5 using maraviroc has been explored in patients with metastatic colorectal cancer (NCT01736813) where 4/6 patients showed a trend towards an increase in CD8^+^ T-cells within the TME (*81*). Also, in this study, the authors identified a partial response (3/5 patients) and stable disease (1/5 patients) when maraviroc was combined with a chemotherapeutic agent (*81*). Although tentative, it highlights intriguing key clinical parallels with our preclinical observations.

In summary, we show that Lyve-1^+^ TAMs derive from an IL-6 polarization program in the TME and demonstrate that the Lyve-1^+^ TAM functions are not always autonomous but can be collaborative through their formation of nests within the Pv space using a CCR5-CCL3/4 axis. We demonstrate that these multi-cellular ‘nest’ structures are biologically important and associated with immune exclusion in the TME and a new resistance mechanism for cytotoxic chemotherapies. This study sheds new light on the collaborative actions of TAMs and suggests their communication could provide novel therapeutic opportunities for targeting their pro-tumoral functions in cancer.

## Acknowledgements

The authors thank Dr Yasmin Haque (KCL) and the NIHR BRC flow cytometry platform at Guy’s and St Thomas’ Biomedical Research Centre for cell sorting and flow cytometry assistance, Dr James Levitt and the Nikon Imaging Centre (KCL) for use of their facilities and assistance with confocal microscopy analyses. The authors would like to thank Drs Paul Lavender, Tracey Mitchell, Gilbert Fruhwirth (KCL) and Professor Awen Gallimore (University of Cardiff) for useful discussion/advice/support, Miss Rosamond Nuamah (KCL) for running the microarray and Mr Stuart Newman (KCL) for help with rederivation of the HO-1^*Luc/eGFP*^ reporter mouse. The authors would like to thank Miss Jiaying Yao for initial optimization of the focal-point macrophage migration assay. This work was funded by a grant from Cancer Research UK (DCRPGF\100009) and the European Research Council (335326). J.N.A is the recipient of a Cancer Research Institute / Wade F.B. Thompson CLIP grant (CRI3645). J.E.A. is supported by the UK Medical Research Council (MR/N013700/1) and is a KCL member of the MRC Doctoral Training Partnership in Biomedical Sciences. F.M.W. is supported by the Medical Research Council (MR/PO18823/1) and Wellcome Trust (206439/Z/17/Z). *MMTV-PyMT* × *Kaede* mice studies were supported by a Cancer Immunology Project Award (C54019/A27535) from Cancer Research UK awarded to D.R.W. The research was supported by the Cancer Research UK King’s Health Partners Centre and Experimental Cancer Medicine Centre at King’s College London, and the National Institute for Health Research (NIHR) Biomedical Research Centre based at Guy’s and St Thomas’ NHS Foundation Trust and King’s College London. The views expressed are those of the authors and not necessarily those of the NHS, the NIHR or the Department of Health.

## Competing interests

Authors declare no competing interests relating to this work.

## Author contributions

J.E.A., J.N.A. conceived the project, designed the approach, performed experiments, interpreted the data wrote the manuscript. J.W.O., I.D., H.P.M., M.B., K.L.A., Z.L., D.C., J.C., D.S., R.B., T.M., W.A., G.L. designed the approach, performed experiments, and interpreted the data. C.E.G., X.Z., F.M.W., T.N., J.M.B., S.K., D.R.W., T.L. designed experiments, interpreted the data and provided key expertise.

## Supplementary Material

### Material and Methods

#### Mice

*MMTV-PyMT* mice used in this study were on an FVB/N background (*28*). Female 4-6 week-old WT Balb/c mice, WT C57BL/6, homozygous *Il6*^-/-^ (B6.129S2-Il6tm1Kopf/J) and *Ccr5*^-/-^ (B6.129P2-Ccr5tm1Kuz/J) and *Lyz2*-cre (B6.129P2-Lyz2tm1(cre)^lfo^/J) C57BL/6 mice were purchased from Charles River. Where indicated, female KO mice were crossed with male *MMTV-PyMT* mice and the F2 homozygous or F2 WT offspring used experimentally. Female C57Bl/6 homozygous *Kaede* mice (*42*) were crossed with male *MMTV-PyMT* (FVB background) mice and the F1 offspring were used experimentally. Cohort sizes were informed by prior studies (*3*, *55*). The *Hmox1^fl/fl^* and *Lyz2* (Lysozyme M) driven cre recombinase were crossed for the *MMTV-PyMT/Hmox1^fl/fl^Lyz2^cre+/-^* (*82*). *Hmox1^fl/fl^* mice were a gift from Professor George Kollias, Biomedical Sciences Research Center “Alexander Fleming”, Athens, Greece. All mice used for experiments were female and randomly assigned to treatment groups. Mice were approximately 21-26 g when tumors became palpable. Experiments were performed in at least duplicate and for spontaneous *MMTV-PyMT* tumor studies individual mice were collected on separate days and all data points are presented. For generation of HO-1-Luciferace-eGFP-knock-in mouse (HO-1^Luc/eGFP^) we have used BAC (Bacterial Artificial Chromosomes) recombineering strategy (*83*). A synthetic cassette containing P2A-Luciferase-P2A-eGFP-Stop-FRT-ßAct-Neo-pA-FRT sequence was inserted before the endogenous stop codon of *Hmox1* in BACs that correspond to *Hmox1* locus. The insert-containing BAC was further subcloned into pR3R4ccdB plasmid that contains gateway sites. The resulting “intermediate” vector contains ~5kb 5’ and 3’ homology arms and was used for generation of the final vector by modular vector assembly by the gateway method. The gateway reaction was performed using LR Clonase II Plus enzyme mix (Thermo Fisher Scientific) according to the manufacturers’ protocol. The intermediate targeting vector was combined with pL3/L4 (DTA selection cassette) and incubated at 25°C overnight (O.N.). After treatment with Proteinase K, the reaction mix was transformed into chemically competent *E. coli* (DH10B, Invitrogen) and plated onto YEG (yeast extract with glucose) agar plates containing 4-chlorophenylalanine and spectinomycin antibiotic (25 μg-mL). Individual colonies were picked and verified with restriction digestion quality control and sequenced across all recombineered junctions. A positive final targeting vector was linearized at *AsisI* restriction site and electroporated to C57BL/6J embryonic stem (ES) cells. Clones were selected and picked under G418 antibiotic selection. Genomic DNA from positively selected ES cell clones were further screened with long range and short-range PCR for target recombination, 5’ and 3’ homology arms using the Sequal Prep Long PCR kit (Thermo Fisher Scientific). The neomycin selection cassette, which was flanked by flippase (Flp) recombination target sites, was removed by fertilizing WT C57BL/6 oocytes with sperm from HO-1^Luc/eGFP^ mice. Six hours after insemination, fertilized zygotes were identified by the presence of pronuclei (87%) and received a cytoplasmic injection of Flp mRNA (Miltenyi Biotech). Injected zygotes were surgically transferred to CD-1 0.5dpc pseudopregnant recipient females. Long range PCR confirmed successful recombination.

#### Cell lines

3B-11 murine endothelial cells and 4T1 mammary adenocarcinoma were obtained from ATCC (*84*). Cell lines were confirmed to be mycoplasma free using the MycoAlert Mycoplasma Detection Kit (Lonza) and were cultured in RPMI (Gibco) supplemented with 10% FCS (Thermo Fisher Scientific).

#### Tumor studies

4T1 (Balb/c) cells were orthotopically implanted for tumors (to generate splenic tumor-derived but TME naive monocytes for *in vitro* studies). A total of 2.5 × 10^5^ cells in 100 μL RPMI were injected subcutaneously into the mammary fat pad of syngeneic female mice. In *MMTV-PyMT* mice, tumors arose spontaneously. When tumors became palpable, volumes were measured every 2-4 days using digital caliper measurements of the long (L) and short (S) dimensions of the tumor. Tumor volume was established using the following equation: Volume= (S^2^xL)/2. *MMTV-PyMT Kaede* mice were photolabeled under anesthesia. Individual tumors were exposed to a violet light (405nm wavelength) through the skin for a total of nine 20 second exposure cycles with a short 5 second break interval between each cycle. Black cardboard was used to shield the rest of the mouse throughout the photoconversion procedure. Mice for 0 h time points were culled immediately after photoconversion. This photoconversion approach was adapted from that used to label peripheral lymph nodes (*85*) and was optimized for *MMTV-PyMT* tumors (*16*). Tumor tissue for flow cytometry analyses was enzyme-digested to release single cells as previously described (*55*, *84*). In brief, tissues were minced using scalpels, and then single cells were liberated by incubation for 60 mins at 37°C with 1 mg/mL Collagenase I from *Clostridium Histolyticum* (Sigma-Aldrich) and 0.1 mg/mL Deoxyribonuclease I (AppliChem) in RPMI. Released cells were then passed through a 70 μm cell strainer prior to staining for flow cytometry analyses. Viable cells were numerated using a hemocytometer with trypan blue (Sigma-Aldrich) exclusion. For drug treatments, drugs were freshly prepared on the day of injection and administered by intraperitoneal (i.p.) injection using a 26 G needle. Maraviroc (Cayman) was solubilized in ethanol and diluted with saline and administered to mice i.p. using a bi-daily dose of 10mg/kg. 5-fluorouracil (Sigma-Aldrich) was prepared fresh and dissolved in saline at 6 mg/mL and injected to mice i.p. at 40 mg/kg/4 days. Immune-depleted mice were injected i.p. every 4 days, starting 48 h prior to the commencement of treatment, with 400 μg of anti-CD8α (53-6.7) (Thermo Fisher Scientific).

#### *In vitro* derived macrophage polarization and gene knockdown

Murine bone marrow (BM) was flushed from the femur and tibia of non tumor bearing WT C57Bl/6 mice using a syringe and needle. Splenocytes for monocyte isolation were acquired from spleens of 4T1 tumor bearing mice by crushing through a 70 μm pore strainer. RBC were lysed using RBC lysis buffer (Roche). Ly6C^+^ monocytes were isolated from the splenocytes by blocking Fc receptors using 5 μg/mL anti-CD16/32 (2.4G2, Tonbo Biosciences) prior to staining with Ly6C PE (HK1.4; Thermo Fisher Scientific) in MACs buffer (DPBS (Thermo Fisher Scientific), 0.5% BSA, 2mM EDTA) at 1 μg/mL, followed by anti-PE MicroBeads (Miltenyi Biotec) and isolated using a MidiMacs separator and LS columns (Miltenyi Biotec) according to the manufacturers’ protocol. BM cells or isolated Ly6C^+^ monocytes were plated in RPMI, 10% FCS, 1 × penicillin/streptomycin (Sigma-Aldrich), 10 ng/mL recombinant murine M-CSF (Bio-Techne) at 1 × 10^6^ cells/well on 6 well plates for 72 h prior to subsequent downstream mRNA and protein analyses. Where viable macrophages were required for ongoing experiments, 5.5 × 10^6^ BM cells were plated at day 0 on 6 cm non tissue culture-treated plates in the above macrophage culture media. Additional murine cytokines, IL-4, IL-6, IFN-γ (Bio-Techne) and/or LPS (Sigma-Aldrich) were added where indicated in the figure legends at 50 ng/mL unless stated otherwise. After 72 h in culture, macrophage purity was assessed by flow cytometry. Macrophages differentiated in the presence of M-CSF alone were referred to as M_(0)_ cells, and macrophages differentiated in the presence of M-CSF and IL-6 were labelled M_(IL-6)_ cells. For siRNA knock down experiments, M_(0)_ macrophages had their media changed to IMDM, 10% FCS and 10 ng/mL M-CSF. In an Eppendorf, ON-TARGET_plus_ SMARTpool siRNA (Horizon Discovery Ltd) targeting *c-Maf* (J-040681-10-0005) *Stat3* (J-040794-10-0005), *Hmox1* (J-040543-12) or ON-TARGET_plus_ non-targeting siRNA (D-001810-01-05), were added to 250 μL of Opti-MEM (Thermo Fisher Scientific) at a concentration of 100 pmol. To each respective tube, an equal volume of Opti-MEM mixed with 5 μL LipofectamineTM RNA_iMAX_ (Thermo Fisher Scientific) was added and incubated for 20 mins at room temperature (RT). The transfection mixture was then drop-wise added to M_(0)_ BMDMs. The wells were gently mixed until the siRNA transfection buffer distributed evenly and incubated for 96 h to allow for protein knock down in the presence or absence of polarizing cytokines at 25 ng/mL as indicated.

#### T-cell isolation

For isolating murine T-cells, spleens were excised from WT C57Bl/6 mice and placed in RPMI, 10% FCS, 20 μM 2-mercaptoethanol, 1X penicillin/streptomycin (Sigma-Aldrich). Spleens were crushed through a 70 μm pore strainer and washed through using RPMI. Liberated splenocytes were centrifuged at 500 × *g* for 3 mins and the cell pellet was re-suspended in 1 mL of red blood cell lysis buffer (Roche) for 2 mins at RT. Cells were then re-centrifuged at 500 × *g* for 3 mins and the pellet was resuspended in RPMI. Live cells were numerated using Trypan blue exclusion on a hemocytometer. CD8^+^ T-cells were purified using the CD8a^+^ T-cell isolation Kit, mouse (Miltenyi Biotec) and isolated using a MidiMacs separator and LS columns (Miltenyi Biotec) according to the manufacturers’ protocol. T-cells were resuspended in T-cell culture media that was further supplemented with 2 ng/mL recombinant murine IL-2 (Bio-Techne) and purified CD8^+^ T-cells were plated at a density of 0.1×10^6^ cells/well in 200 μL onto a high binding 96-well plate (Sigma-Aldrich) that had been pre-coated O.N. with a mix of anti-mouse CD3ε (145-2C11, 5 μg/mL) and anti-mouse CD28 (37.51, 3 μg/mL) antibodies in sterile DPBS (100 μL/well) at 4°C. After 48 h CD8^+^ T-cells were transferred to a fresh uncoated plate and rested for at least 48 h before being numerated and used for down-stream *in vitro* assays.

#### *In vitro* macrophage focal-point migration assay

M_(0)_ macrophages were generated as described above and removed from the plate using 1 mL of enzyme-free dissociation media (Thermo Fisher Scientific) and mechanically detaching the cells using a cell scrapper. The resultant solutions were centrifuged at 2000 *x g* for 3 mins and resuspended at a concentration of 10,000 cells/mL in 80% RPMI (inc.10%FCS) and 20% methylcellulose with 10 ng/mL M-CSF. To prepare a methylcellulose stock solution; autoclaved methylcellulose (Sigma-Aldrich) was dissolved at 24 g/L in pre-heated serum-free RPMI media for 2 mins at 60°C. After this, the solution was diluted with 2 volumes of RT serum-free RPMI and then mixed O.N. at 4°C. The final solution was cleared by centrifugation at 5000 *x g* for 2 h at RT. To the lid of an inverted petri dish, 25 μL drops of macrophages were placed, the lid was carefully inverted and placed on top of a DPBS filled petri dish. The dish was incubated for 24 h to allow the formation of macrophage spheroids within the hanging drop. Subsequently, media containing spheroids were collected in Eppendorf tubes using DPBS. The spheroids were allowed to briefly settle in the bottom of the Eppendorf tubes after which the DPBS was carefully removed and the spheroids were placed in wells containing RPMI, 10% FCS, 10 ng/mL M/CSF with or without 25 ng/mL IL-6 and placed in a 37°C 5% CO_2_ incubator to allow attachment and eventual spreading. Images of spheroid cultures was performed using the live cell Eclipse Ti-2 inverted microscope in the Nikon Imaging Centre at King’s College London. NIS Elements Advanced Research software (Nikon) was used to process the images. Total area of spheroids was measured using the “Analyze Spheroid Cell Invasion In 3D Matrix” macro in ImageJ, and cell number was counted using the “Cell counter” plugin in ImageJ.

#### *In vitro* perivascular nest transwell assay

Transwell assays were conducted with transwell inserts with 8 μm pores (Corning) for migration studies and 0.4 μm pores (Corning) for permeability studies. Inserts were coated with Basement Membrane Extract (Cultrex) diluted 1:100 in RPMI for 1 h at RT. Excess Basement Membrane Extract was aspirated and 2 × 10^4^ 3B-11 endothelial cells were seeded onto the apical side of the transwell insert in RPMI supplemented with 10% FCS and left to attach for 24 h. Media was removed, the whole plate inverted and 10^5^ M_(0)_ or M_(IL-6)_ BMDMs were seeded onto the basolateral side of the transwell membrane in RPMI supplemented with 10% FCS and left to attach for 2 h at 37°C. Subsequently, the plate was reinverted to its original position and RPMI supplemented with 10% FCS, 10 ng/mL M-CSF with or without 10 ng/mL IL-6 added to the apical and basolateral space. After cells were left to interact for 24 h at 37°C, 4 × 10^5^ CD8^+^ T cells (which had been prior incubated on anti-CD3 and -CD28 coated plated to develop effector function) were added to the inserts in RPMI, 10% FCS and 100 ng/mL with murine CXCL10 (Bio-Techne) spiked into the wells. After 16 h, migrated cells were collected from the well and stained for flow cytometry analysis and quantification with AccuCheck counting beads (Thermo Fisher Scientific). The permeability assay was performed as described previously (*86*). In brief, permeability of the *in vitro* perivascular niche was measured using 4% (w/v) Evans Blue-conjugated Bovine Serum Albumin (BSA) diluted in DPBS was placed into the apical transwell chamber while phenol red-free RPMI, 10% FCS was added to the basolateral chamber. Presence of Evans Blue-BSA in the basolateral chamber was assessed at indicated time-points by absorbance at 620 nm on a NanoDrop^™^ spectrophotometer (Thermo Fisher Scientific). These experiments were performed in the presence or absence of 10 ng/mL IL-6 and M-CSF, 25 μM Sn (IV) mesoporphyrin IX dichloride (SnMP: Frontier Scientific). SnMP was prepared fresh on the day and dissolved as previously described (*55*).

#### Western blot

Cells were lysed and SDS PAGE/western blots were conducted as previously described (*3*). In brief, cells were lysed in the well using Western blot lysis buffer 0.1M Tris-hydrochloride pH 6.8, with 20% glycerol and 4% sodium dodecyl sulphate containing 1X protease and phosphatase inhibitor cocktail (Thermo Fisher Scientific). All tubes were heated at 95°C for 15 mins to break down DNA. Protein concentration was then determined using the PierceTM BCA Protein Assay Kit (Thermo Fisher Scientific) using the manufacturers’ protocol. Samples were then run under reducing conditions on 12% bis-tris sodium dodecyl sulphate polyacrylamide gel electrophoresis (SDS-PAGE) gels alongside SeeBlue^™^ Plus2 pre-stained makers (Thermo Fisher Scientific). SDS-PAGE gels were then transferred onto polyvinyl-difluoride (PVDF) membranes which were subsequently blocked in 100 mM Tris, 140 mM NaCl, 0.1% Tween 20, pH7.4 (TBS-T) containing 5% skimmed milk at RT for 1 h. Primary antibodies were applied at 4°C O.N. and secondary antibodies for 1 h at RT. Wash steps to remove unbound antibodies were 3 × 20 mins in TBS-T. The following primary antibodies were used: rabbit anti-β-actin, 1:5,000 (ab8227, Abcam), rabbit anti-HO-1 1:1,000 (10701-1-AP, Proteintech), rabbit anti-c-MAF 1:1,000 (ab77071, Abcam), rabbit anti-STAT3 1:2000 (79D7, Cell Signalling). These antibodies were detected using goat anti-rabbit immunoglobulins/HRP secondary antibody 1:2,000 (Agilent Dako). Then, protein bands were detected using Luminata^™^ Crescendo Western HRP substrate (Millipore) and CL-XPosure^™^ Film (Thermo Fisher Scientific).

#### Immunofluorescence

Mouse mammary tumor tissue or human invasive ductal carcinoma tissue was fixed O.N. in 4% paraformaldehyde, followed by O.N. dehydration in 30% sucrose prior to embedding in OCT and snap freezing absolute ethanol and dry ice. Sections from the embedded tumors (10 μm) were placed onto microscope slides were incubated further in 4% paraformaldehyde in DPBS for 10 mins at RT prior to washing in TBS-T and blocked using TBS-T, 10% donkey serum (Sigma-Aldrich), 0.2% Triton X-100. Immunofluorescence staining was performed as previously described (*3*). Antibodies against the following targets and their dilutions were used as follows; αSMA 1:100 (AS-29553, Anaspec), CD31 1:100 (ab28364 Abcam), CD31 1:100 (MA1-40074, Thermo Fisher Scientific), CD31 1:100 (EP3095, Abcam), CD68 1:100 (KP1, Invitrogen), F4/80 1:100 (C1:A3-1, Bio-RAD), HO-1 1:100 (AF3776, R&D systems), HO-1 1:100 (10701-1-AP, Proteintech), Lyve-1 1:100 (ab33682, Abcam). Primary antibodies were detected using antigen specific donkey IgG, used at 1:200: AlexaFluor^®^ 488 anti-rabbit IgG, AlexaFluor^®^ 488 anti-rat IgG, AlexaFluor^®^ 488 anti-goat IgG, AlexaFluor^®^ 568 anti-rabbit IgG, AlexaFluor^®^ 568 anti-goat IgG, AlexaFluor^®^ 647 anti-rabbit IgG (Thermo Fisher Scientific), AlexaFluor^®^ 647 anti-mouse IgG, NL637 anti-rat goat IgG (R&D Systems) and Cy3 anti-sheep IgG (Jackson ImmunoResearch). Viable blood vessels were visualized in mice through intravenous (i.v.) injection of FITC-conjugated dextran (2,000,000 MW, Thermo Fisher Scientific) 20 mins prior to sacrifice. Nuclei were stained using 1.25 μg/mL 4’,6-diamidino-2-phenylindole,dihydrochloride (DAPI) (Thermo Fisher Scientific). RNA scope was performed on formalin-fixed paraffin-embedded (FFPE) *MMTV-PyMT* tumor sections as per manufacturers’ instructions using the RNAscope^®^ Multiplex Fluorescent Reagent Kit v2 Assay (Bio-Techne; 323100-USM). The Mm-IL6 (Bio-Techne; 315898) probe was used and was detected using the Opal^™^ 570 Reagent Pack (FP1488001KT, Akoya Biosciences). Following RNAscope, immunofluorescence imaging was performed as previously described above. Images were acquired using a Nikon Eclipse Ti-E Inverted spinning disk confocal. For 5-color confocal microscopy, images were acquired on a Nikon A1R spectral deconvolution confocal microscope. Using a 32-channel A1-GasAsP-detector unit, fluorochrome emission can be split up in up to 32 bands from 400 to 750 nm with a spectral discrimination of 10 or 20 nm bandwidth when excited by a new laser-box with 4 solid state lasers 405, 488, 561 and 640 nm. The acquisition signals were then clearly and reliably distinguished by a process called “spectral unmixing”. Images were analyzed using the NIS-Elements software.

#### Bioluminescence imaging

For assessing Luc bio-distribution *in vivo* mice were injected i.p. with 3 mg XenoLight D-luciferin (PerkinElmer) in sterile DPBS 10 mins prior to imaging. For whole-body imaging animals were anesthetized and placed in the *in vivo* Imaging System (IVIS^®^) Lumina Series III (PerkinElmer). For imaging the Luc bio-distribution of different tissues, the mice were injected with D-luciferin and sacrificed after 10 mins and the dissected tissues were then imaged 15 mins later. To quantify luminescence, a region of interest (ROI) was drawn around a specific area and total photon flux (PF) (photon/second; p/s) was measured. All data was analyzed using the Living Image Software (PerkinElmer).

#### Flow cytometry

Flow cytometry was performed as previously described (*32*). The following antibodies against the indicated antigen were purchased from Thermo Fisher Scientific and were used at 1 μg/mL unless stated otherwise: CCR5 PE (HM-CCR5(7A4)), CD3ε APC, PE (145-2C11) and BV421 (17A2; Biolegend^®^), CD4 APC and FITC (RM4-5), CD8α BV421 and FITC (53-6.7; Biolegend^®^), CD8β FITC and eFluor^®^450 (H35-17.2), CD11b BV510 (M1/70; Biolegend^®^), CD11c APC (N418) and FITC (N418; Biolegend^®^), CD16/32 (2.4G2; Tonbo Biosciences), CD19 BV421, APC (6D5; Biolegend^®^) and FITC (1D3/CD19; Biolegend^®^), CD31 FITC, PE (390) and BV510 (MEC 13.3; BD biosciences), CD45 BV605, APC, APC-eFluor^®^ 780 (30-F11) and BV510 (30-F11; Biolegend^®^), CD90.1 eFluor^®^ 450 (HIS51) and BV510 (OX-7; Biolegend^®^), CD90.2 eFluor^®^ 450 (53-2.1) and BV510 (53-2.1; Biolegend^®^), CD206 APC, FITC and BV786 (C068C2; Biolegend^®^) and APC (FAB2535A; Bio-Techne), F4/80 APC, APC-eFluor^®^ 780, PE and eFluor^®^ 660 (BM8), BV421 and FITC (BM8; Biolegend^®^), Foxp3 PE-Cyanine5 (FJK-16s), Gr-1 FITC (RB6-8C5; Biolegend^®^), Granzyme-B PE (GB11), ICAM-1 BV421 (YN1/1.7.4, Biolegend^®^), IFN-γ APC (XMG1.2), Ly6C APC, APC-eFluor^®^ 780, eFluor^®^450 (HK1.4) and FITC (HK1.4; Biolegend^®^), Ly6G (1A8; Biolegend^®^), Lyve-1 Alexa Fluor^®^ 488, PE (ALY7) and APC (FAB2125A; Bio-Techne), MHCII PE (M5/114.15.2), BV421, BV510 and FITC (M5/114.15.2; Biolegend^®^), NK1.1 APC (PK136), pNAD AF647 (MECA-79, Biolegend^®^), TNF-α PE (MP6-XT22), VCAM-1 PE (429 vCAM.A, Biolegend^®^). Positive stains were compared to fluorescence minus one (FMO) controls. Intracellular stains were performed as previously described (*32*). Dead cells and red blood cells were excluded using 1 μg/mL 7-amino actinomycin D (7AAD; Sigma-Aldrich), Fixable Viability Dye eFluor^®^ 780 or Near-IR Dead cell staining kit (Thermo Fisher Scientific) or DAPI alongside anti-Ter-119 PerCP-Cy5.5 (Ter-119). Data were collected on a BD FACS Canto II (BD Biosciences). Cells were sorted on a BD FACSAria (BD biosciences). Data was analyzed using FlowJo software (BD biosciences). Immune cells (CD45^+^) were separated based upon the following surface characteristics: CD11c^+^F4/80^-^ (dendritic cells), CD11b^+^F4/80^hi^ (macrophages), F4/80^-/lo^Ly6G’Ly6C^+^ (monocytes), CD11b^+^Ly6G^+^ (neutrophils), NK1.1^+^ (NK/NKT-cells), CD3ε^+^ (T-cells), CD3ε^+^CD4^+^ (CD4^+^ T-cells), CD3ε^+^CD4^+^Foxp3^+^ (Tregs cells), CD3ε^+^CD8α/β^+^ (CD8^+^ T-cells), CD3ε^-^CD19^+^ (B cells). Cancer associated fibroblasts (CAFs) were identified as CD45^-^ Thy1^+^ cells and tumor cells were identified as CD45^-^Thy1^-^CD31^-^.

#### Quantitative real time PCR

mRNA was extracted and quantitative reverse transcription PCR was performed as previously described (*32*) using the following primers/probes purchased from (Thermo Fisher Scientific): *Il6* Mm00446190_m1 and *Tbp* Mm01277045_m1. Expression is represented relative to the house-keeping gene Tata-binding protein (*Tbp*). Gene expression was measured using an ABI 7900HT Fast Real Time PCR instrument (Thermo Fisher Scientific).

#### Transcriptomic data and analysis

TAM, CAF and endothelial Bulk RNAseq and human and mouse TAM scRNA-seq datasets were previously published and described (*16*, *29*) and datasets are publicly accessible (see ‘Data Availability’ section). Downstream analysis was performed using the Seurat v3 R package (*87*) and analysis pipeline outlined in (*16*). For upstream regulator analysis we used the QIAGEN IPA (QIAGEN Inc., https://digitalinsights.qiagen.com/IPA) (*43*). When comparing scRNA-seq datasets between human and mouse TAMs, the Garnett package was used (*88*) which has previously been employed to perform mouse-human cross-species comparative analysis (*89*). Murine data and a marker file specifying LYVE1 were provided to Garnett and the model was trained (train_cell_classifier()) with default settings, using the same 2000 genes with highest variance chosen for clustering previously (*16*). Publicly available human data (*29*) were then classified (classify_cells()) with default settings.

Results were projected and plotted on the associated UMAP coordinates from the same data using a customised R script. For Illumina microarray analysis purified mRNA for the respective polarized splenocyte-derived macrophages were cultured for isolated using the PureLink^®^ RNA Mini Kit (Ambion) according to the manufacturers’ protocol. The purity of the isolated mRNA was assessed using a NanoDrop^™^ spectrophotometer (Thermo Fisher Scientific) and the quality and integrity using an Agilent 2100 Bioanalyzer (Agilent Technologies). mRNA was converted to cDNA, then subsequently amplified using the Ovation^®^ PicoSL WTA system V2 (NuGen), biotinylated using the Encore^®^ BiotinIL Module (NuGen), and then hybridized to MouseWG-6 V2.0 Beadchip microarray (Illumina).

Following hybridization, the arrays were washed, blocked, and stained with streptavidin-Cy3 using the Whole-Genome Gene Expression Direct Hybridisation Assay (Illumina). Microarrays were run on an Illumina iScan system, raw fluorescence signals were collected using GenomeStudio (Illumina), and the data imported into Partek Genomics Suite for analysis. Background was subtracted from the raw data and fluorescence signals were normalized using the quantiles method (*90*). All p-values were adjusted for multiple testing using the procedure of Benjamini and Hochberg.

#### Statistics

Normality and homogeneity of variance were determined using a Shapiro-Wilk normality test and an F-test respectively. Statistical significance was then determined using a two-sided unpaired Students *t* test for parametric, or Mann-Whitney test for nonparametric data using GraphPad Prism 8 software. A Welch’s correction was applied when comparing groups with unequal variances. For microarray gene analysis, significance of differences (fold change) between the groups were assessed with Partek^®^ Genomics Suite^®^ software (Partek^®^) using an ANOVA test. Correction for multiple hypotheses was applied to *P*-values by controlling the percentage of false discovery rate. Adjusted *P*-values of < 0.01 were considered significant. Statistical analysis of tumor growth curves was performed using the “CompareGrowthCurves” function of the statmod software package (*91*). No outliers were excluded from any data presented.

#### Study approval

All experiments involving animals were approved by the Animal and Welfare and Ethical Review Board of King’s College London or the University of Birmingham and the Home Office UK. Human breast adenocarcinoma tissue was obtained with informed consent under ethical approval from the King’s Health Partners Cancer Biobank (REC reference 12/EE/0493).

#### Data availability

The RNA-seq transcriptomic and microarray datasets that support the findings of this study are available through the Gene Expression Omnibus; GSE160561, GSE160641, GSE113034. The microarray datasets are available at GSE192911. The authors declare that all other data supporting the findings of this study are available within the paper and its supplementary information files.

### Supplementary Figures

**Table S1.** Cellular classification of the ‘murine Lyve-1^+^ TAM-like cells’ in the human breast cancer scRNA-seq dataset. Table shows the cell classification for Lyve-1^+^ TAM-like cells within the human breast cancer scRNA-seq atlas (*29*). Lyve-1^+^ TAM-like cells were predominately associated as a subset of cells within the monocyte/macrophage classification (1,342/1,730). Cell classifications are taken from (*29*).

| **Cell classification** | **Count** |
| --- | --- |
| B-cell Memory | 10 |
| CAFs MSC iCAF-like | 4 |
| CAFs myCAF-like | 3 |
| Cancer Basal SC | 6 |
| Cancer Cycling | 2 |
| Cancer LumA SC | 1 |
| Cycling PVL | 1 |
| Cycling Myeloid | 99 |
| DCs | 4 |
| Endothelial ACKR1 | 94 |
| Endothelial CXCL12 | 9 |
| Endothelial Lymphatic LYVE1 | 58 |
| Endothelial RGS5 | 50 |
| Macrophage | 1181 |
| Mature Luminal | 1 |
| Monocyte | 160 |
| Myoepithelial | 2 |
| NK cells | 1 |
| Plasmablasts | 4 |
| PVL Differentiated | 17 |
| PVL Immature | 18 |
| T-cells CD4<sup>+</sup> | 3 |
| T-cells CD8<sup>+</sup> | 2 |
| **Total** | **1730** |

**Figure S1.**
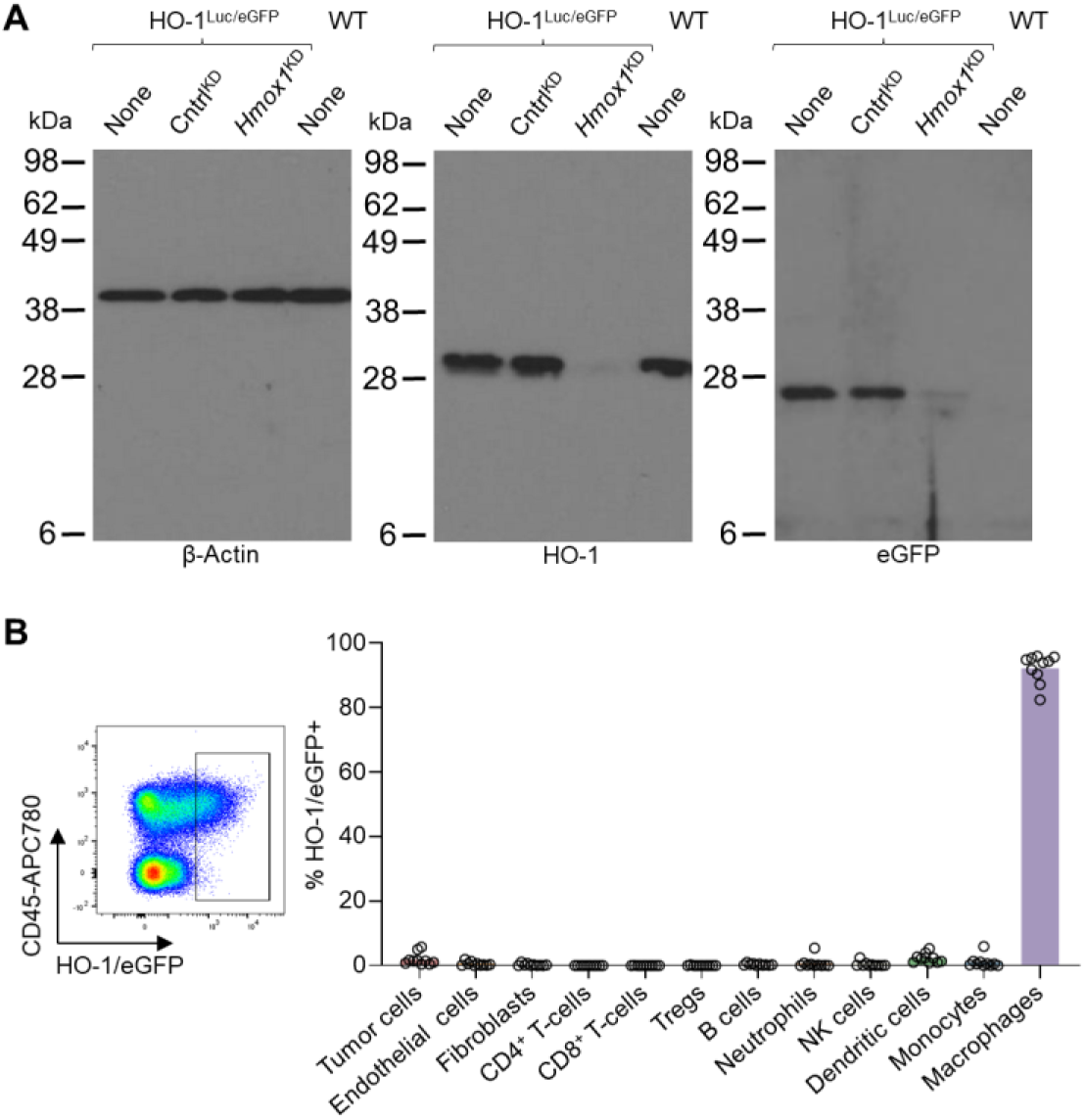
Validation of HO-1^Luc/eGFP^ reporter and cellular restriction of HO-1 expression in the TME. (**A**) Full western blot images of those displayed in Fig. 1L for probing the indicated proteins in the conditions with none,Cntrl (control) and *Hmox1* knockdown (KD) conditions in BMDMs from HO-1^Luc/eG^FP and WT mice as indicated. (**B**) Representative flow cytometry plot of FACs-gated live (7AAD^-^) HO-1/eGFP expressing cells from enzyme-dispersed tumors from MMTV-PyMT HO-1^Luc/eGFP^ mice. Positive cells gated based on FMO control (left panel). Phenotype of the gated HO-1/eGFP expressing cells in the TME (n=10 mice). Markers used to differentiate the individual cell type are described in Material and Methods. Bar charts represent mean and the dots show individual data points from individual tumors and mice.

**Figure S2.**
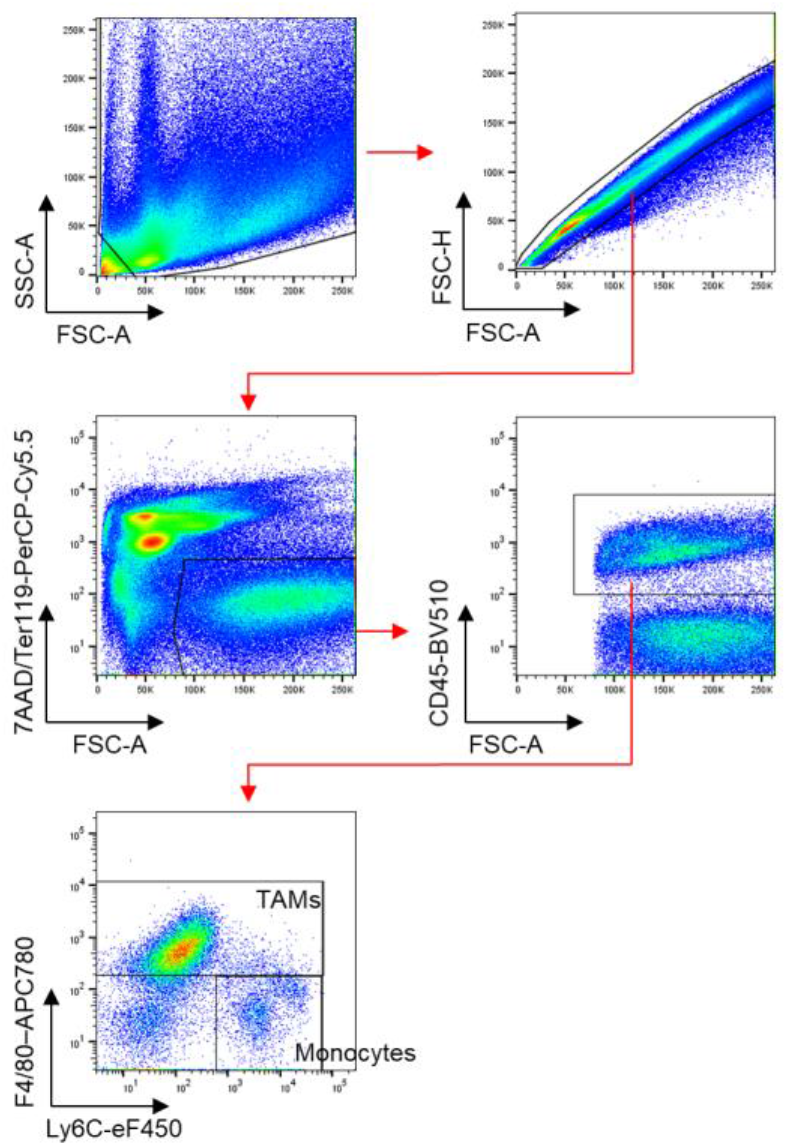
Flow cytometry gating strategy of TAMs to analyze HO-1/eGFP expression. Representative gating strategy for identifying live (7AAD^-^) TAM populations in enzyme-dispersed tumors from *MMTV-PyMT* HO-1^Luc/eGFP^ mice stained for the markers shown and identified using flow cytometry.

**Figure S3.**
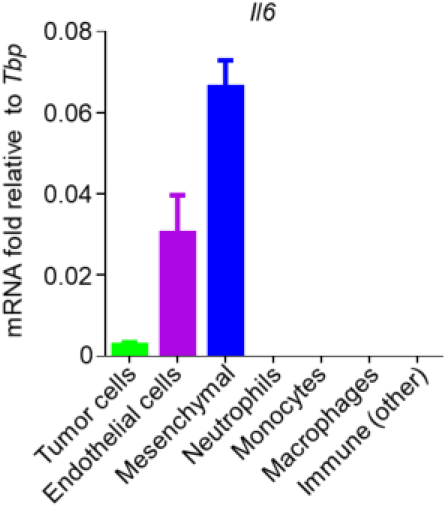
Tumoral *Il6* mRNA expression. The indicated tumoral populations were sorted using flow cytometry from an enzyme-dispersed *MMTV-PyMT* tumor, bars represent the mean and error bar showing the s.d. between wells (representative of duplicate experiments). Markers used for gating can be found in Material and Methods section.

**Figure S4.**
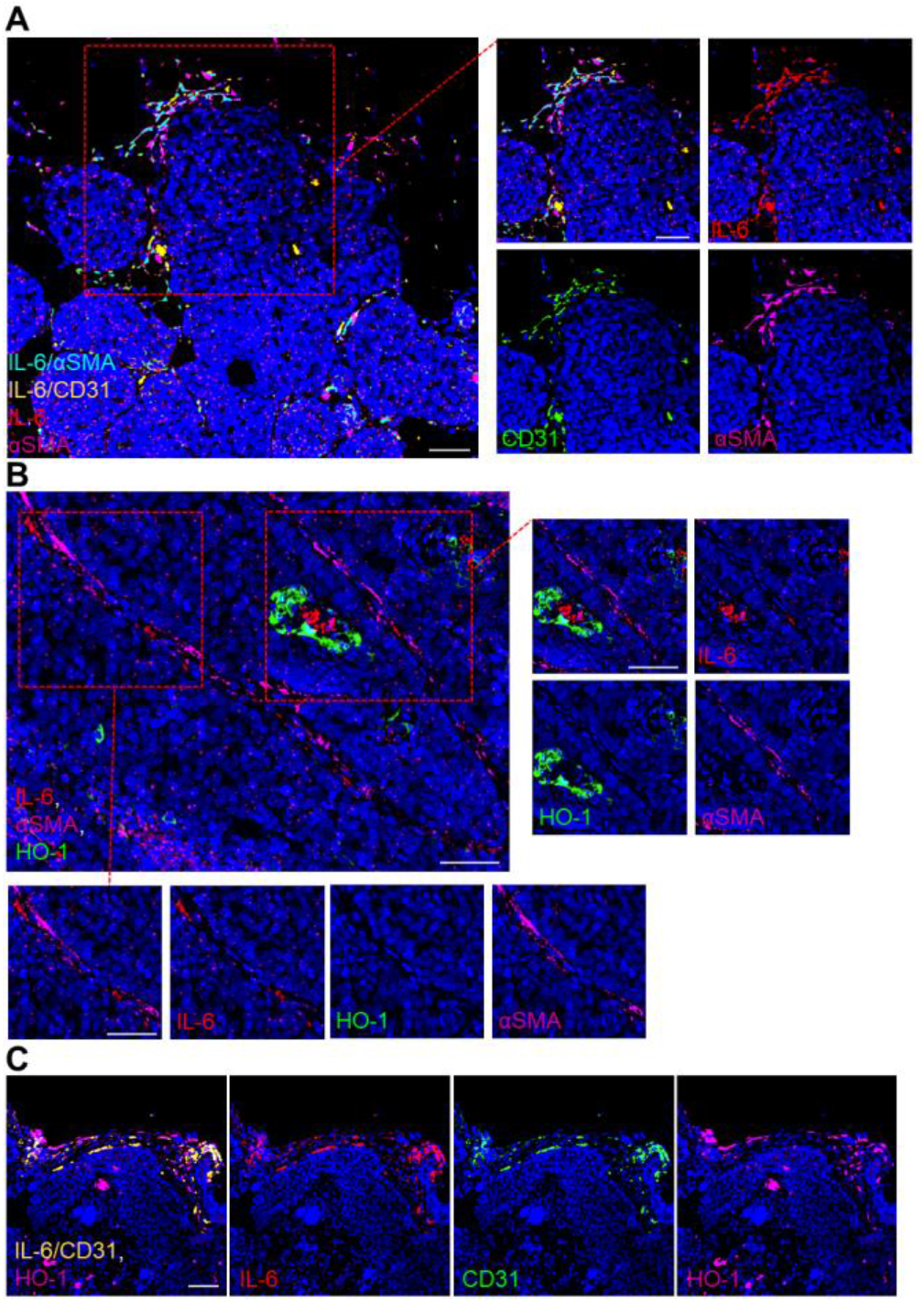
CD31 expressing cells express *Il6* mRNA proximal to HO-1^+^ TAMs in the Pv niche of *MMTV-PyMT* tumors. Representative images of a FFPE section from a *MMTV-PyMT* tumors. (**A**) Tumor sections stained with DAPI (nuclei;blue) and antibodies against again CD31 (green), αSMA (magenta) and probed for *ll6* mRNA (red). Co-localization of *ll6* mRNA and αSMA is displayed in cyan and co-localization of *ll6* mRNA and CD31 is displayed in yellow. (**B**) Tumor sections stained with DAPI (nuclei;blue) and antibodies against again HO-1 (green), αSMA (magenta) and probed for *ll6* mRNA (red). (**C**) Tumor sections stained with DAPI (nuclei; blue) and antibodies against again CD31 (green), HO-1 (magenta) and probed for *ll6* mRNA (red). Co-localization of *ll6* mRNA and CD31 is displayed in yellow. Scale bars in Figure represent 50 μm.

**Figure S5.**
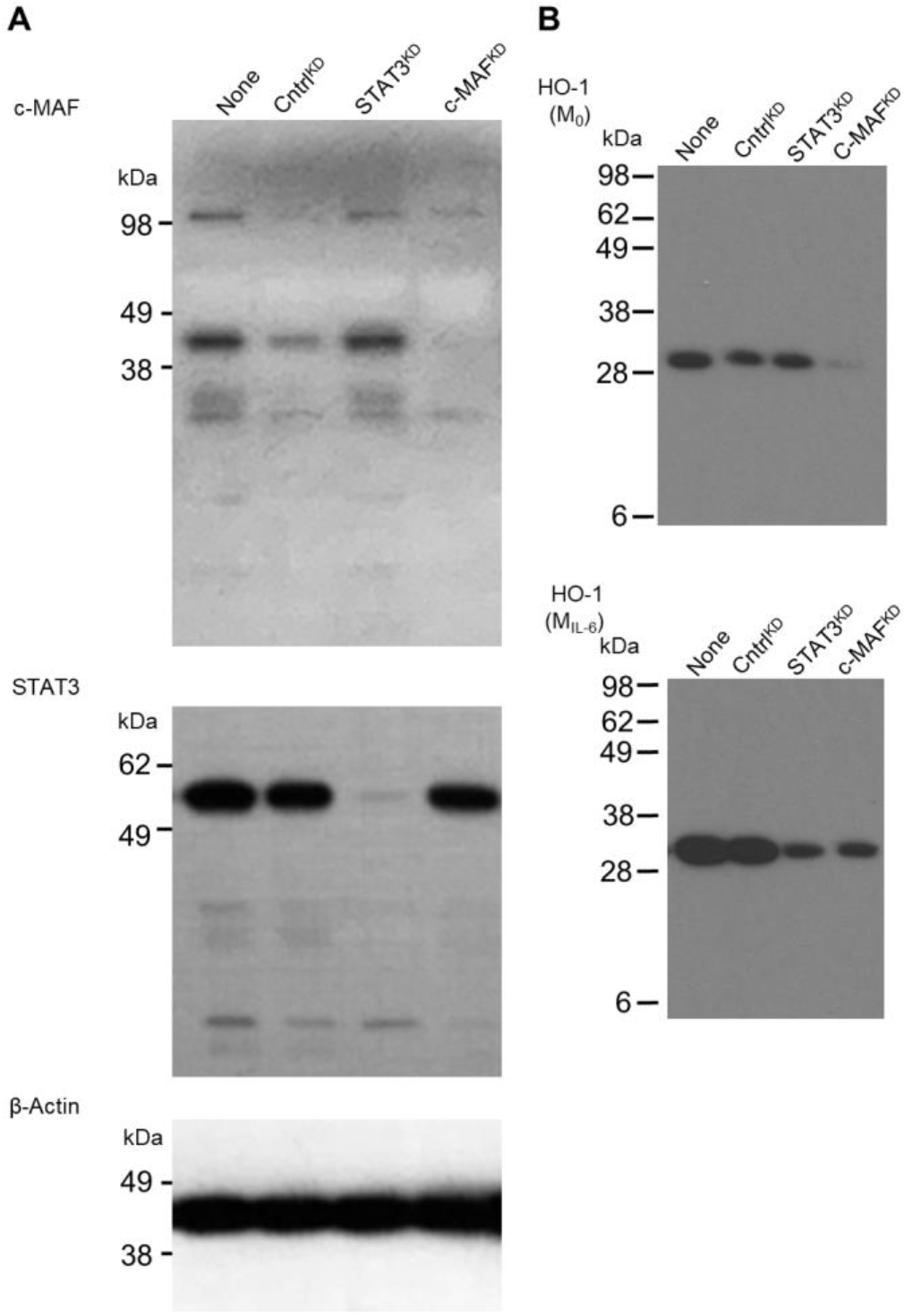
Full western blot images of c-MAF, STAT3 and HO-1 in IL-6 polarized BMDMs. (**A-B**) Full western blot images of those displayed in Fig. 3I probed for STAT3 and c-MAF (**A**) and HO-1 (**B**) expression in lysates from BMDMs derived from Cntrl (control), STAT3 and c-MAF knockdown conditions.

**Figure S6.**
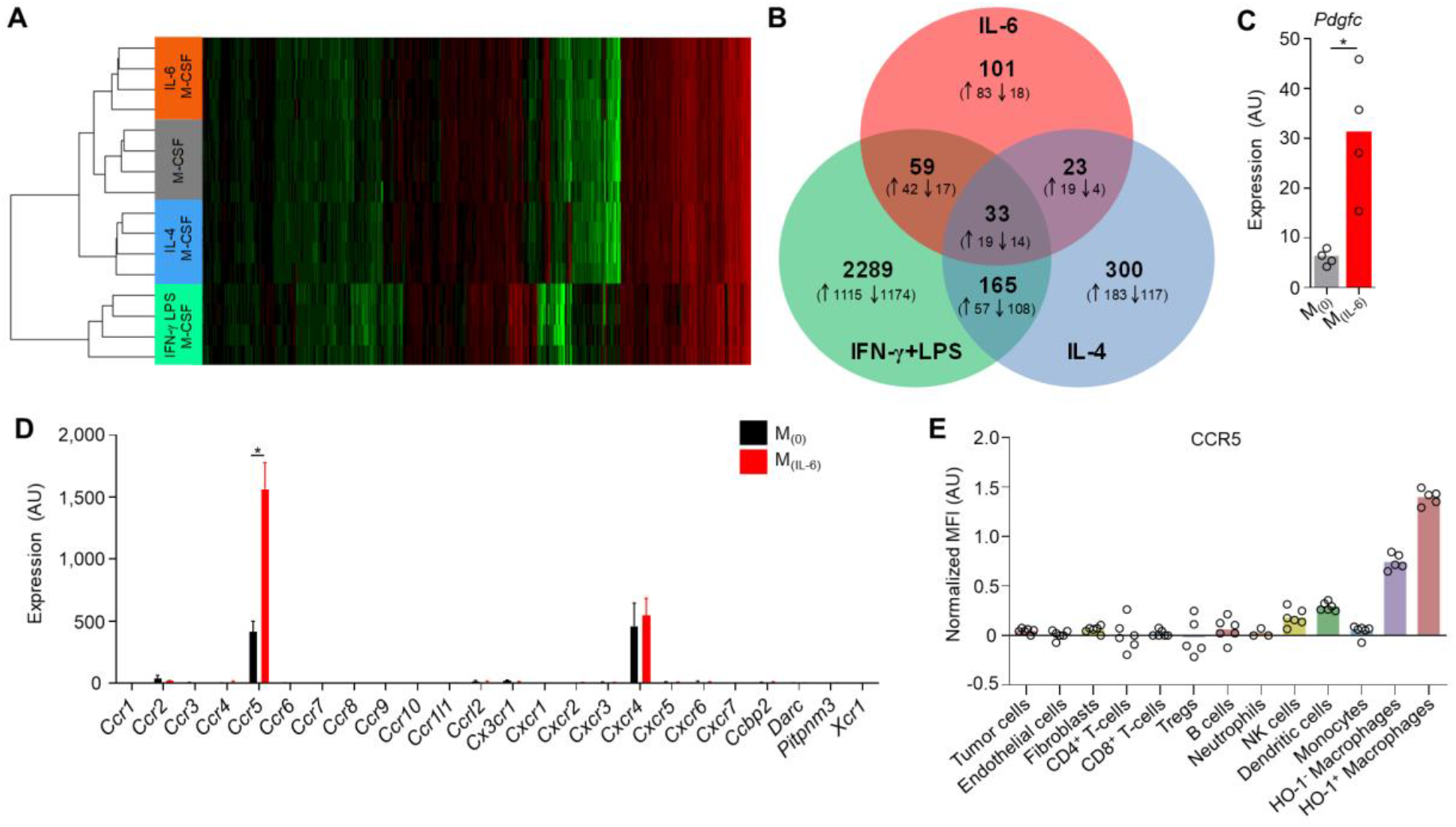
Macrophage polarization by IL-6 induces CCR5 expression. (**A-D**) Splenic monocyte-derived macrophages (as described in Material and Methods) were exposed for 72 h to M-CSF alone (M_(0)_) or M-CSF and IL-6 (M_(IL-6)_) or IL-4 (M1) or IFN-γ/LPS (M2). mRNA was extracted and analyzed using a microarray (n=4 per condition). (**A**) Heatmap of gene expression. (**B**) Venn diagram numerating the differentially regulated genes (up arrow denotes up-regulated and down arrow denotes down-regulated) between the polarization states of the macrophages. Relative expression of *Pdgfc* (**C**) and a range of chemokine receptors (**D**) between M_(0)_ and M_(IL6)_ macrophages. (**E**) Normalized (to total TAMs) median fluorescent intensity (MFI) of the surface expression of CCR5 on the indicated cell populations against fluorescence minus one (FMO) staining using flow cytometry analysis from enzyme-dispersed *MMTV-PyMT* HO-1^Luc/eGFP^ tumors where HO-1 expressing TAMs were gated based upon the presence of absence of HO-1/eGFP. Bar charts represent the mean and the dots show individual data points from individual tumors and mice, error bars represent s.d. * *P*<0.05.

**Figure S7.**
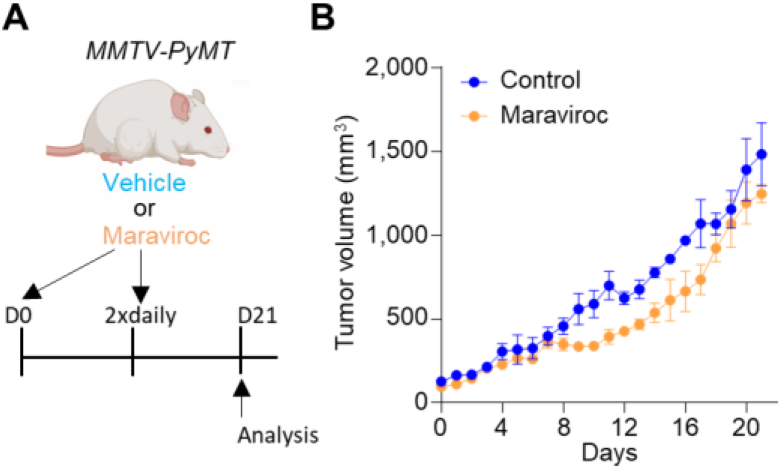
The CCR5 inhibitor maraviroc does not affect tumor growth in *MMTV-PyMT* tumors. (**A**) Schematic representing the dosing strategy for the CCR5 inhibitor maraviroc (bi-daily dose of 10 mg/kg) and vehicle in *MMTV-PyMT* mice (n=3 per group). (**B**) Growth curves of *MMTV-PyMT* tumors in mice treated with vehicle or maraviroc as shown in (**A**) (n=3 mice per group). Line charts display the mean and SEM.

**Figure S8.**
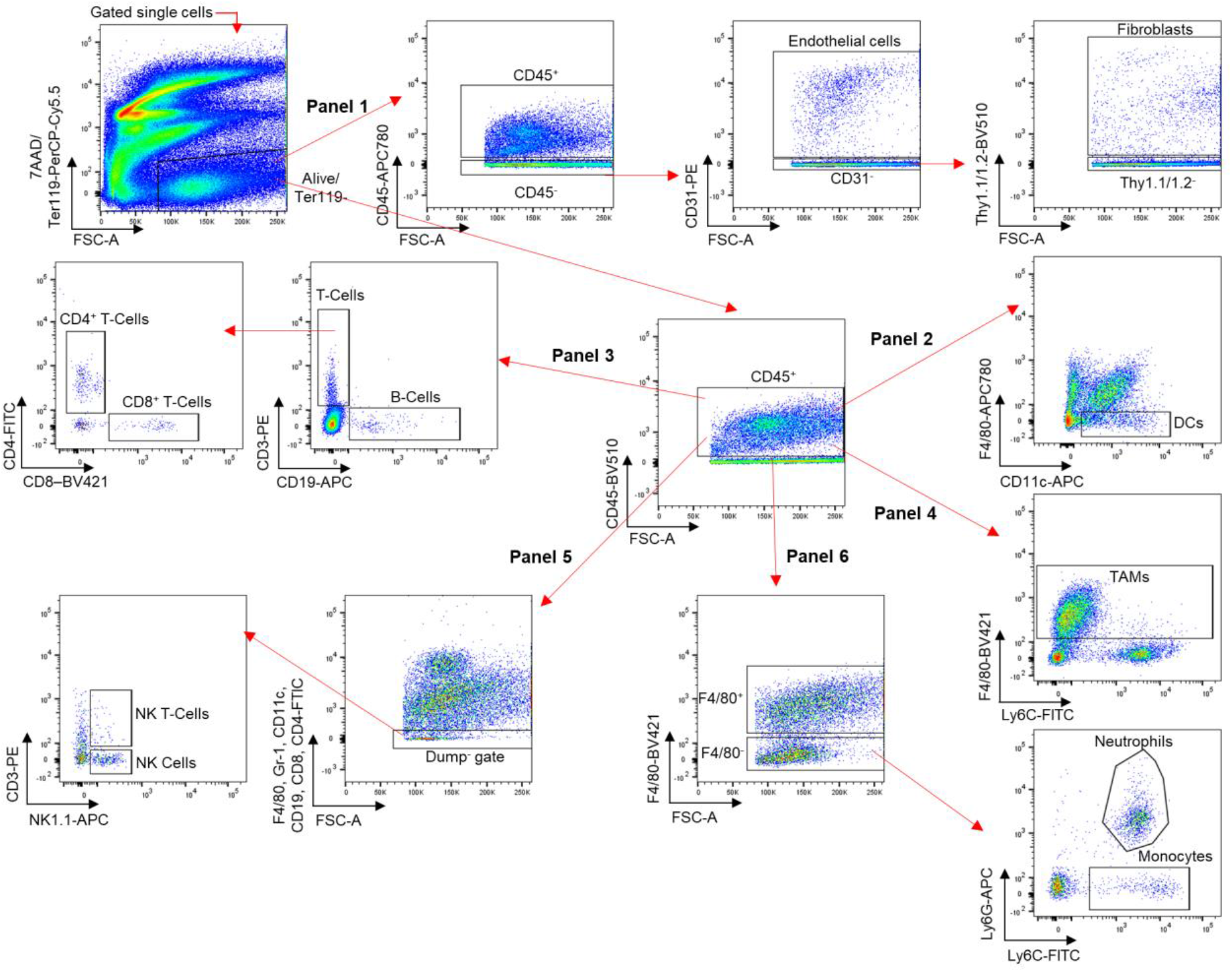
Flow cytometry gating strategy for stromal populations. Representative gating strategy for identifying live (7AAD^-^) nucleated (Ter119^-^) stromal populations in enzyme-dispersed tumors from *MMTV-PyMT* mice using flow cytometry. Positive gates are applied based upon FMO stains.

**Figure S9.**
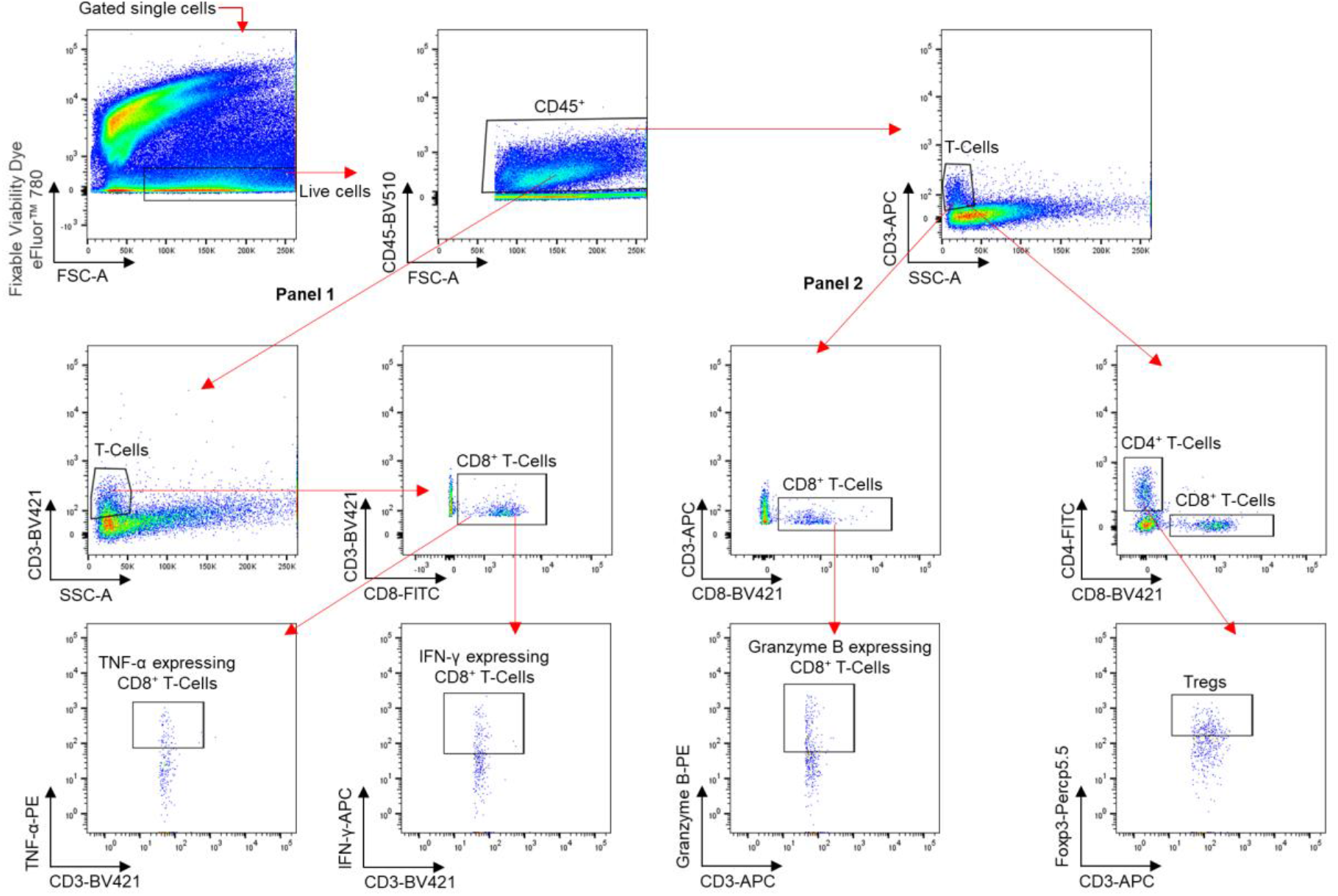
Flow cytometry gating strategy for Foxp3^+^ Tregs and T-cell cytokines. Representative gating strategy for identifying live (fixable viability dye^-^) T-cell populations in enzyme-dispersed tumors from *MMTV-PyMT* mice using flow cytometry. Positive gates are applied based upon FMO stains.

**Figure S10.**
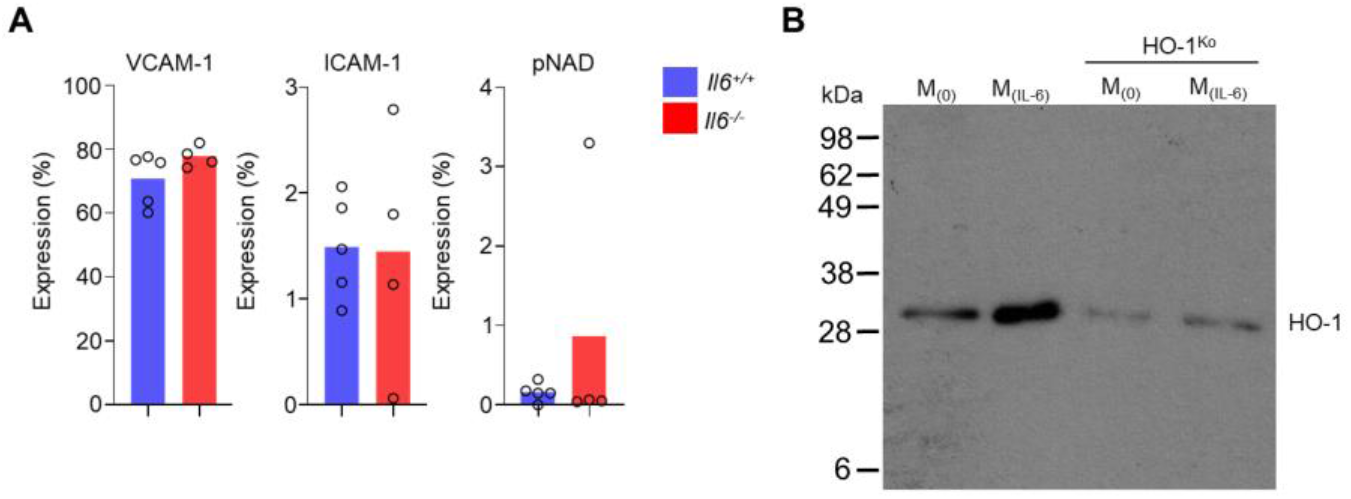
Vascular adhesion molecule expression in the TME and full HO-1^KO^ western. (**A**) Abundance of live (7AAD^-^) CD45^-^ CD31^+^ endothelial cells in enzyme-dispersed tumors from *Il6*^+/+^ and *Il6*^-/-^ *MMTV-PyMT* mice assessed using flow cytometry for expression of the indicated markers (n=4-5 per group). (**B**) Full western blot images of those displayed in Fig. 5H probing for HO-1 expression in BMDMs from mice carrying the *Hmox1*^fl/fl^ allele with (right two columns) and without (left two columns) *Lyz2*^cre^ in M_(0)_ (M-MCF alone) and IL-6 polarization (M_(IL-6)_) conditions. Bar charts represent the mean and the dots show individual data points from individual tumors and mice.

**Figure S11.**
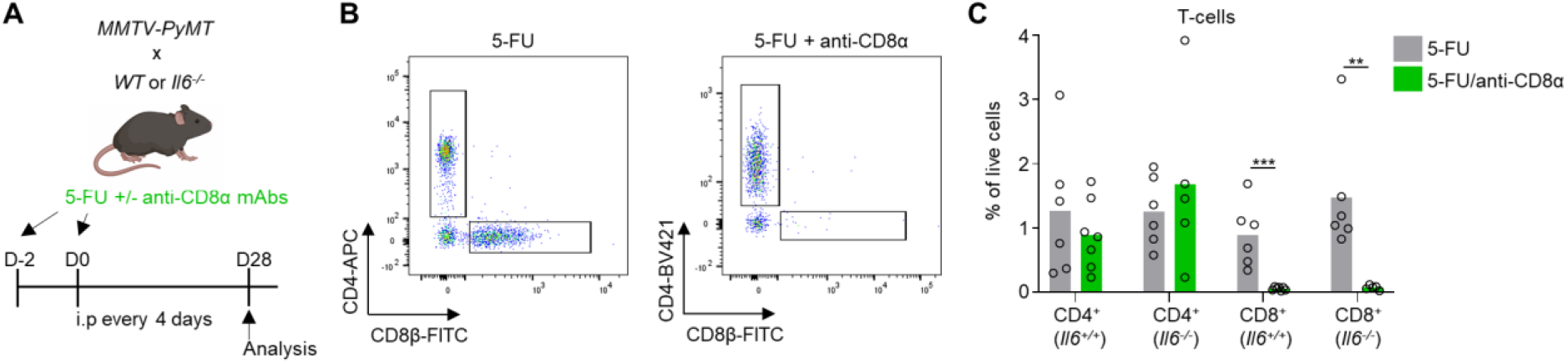
Depletion of tumor infiltrating CD8^+^ T-cells in *MMTV-PyMT* mice. (**A**) Schematic representing the dosing strategy for the immune-depleting anti-CD8α antibodies in *MMTV-PyMT* mice shown in Fig. 5J. (**B**) Flow cytometry gating strategy for tumor infiltrating CD8^+^ (CD3^+^ CD8β^+^) and CD4^+^ (CD3^+^ CD4^+^) T-cells from a *MMTV-PyMT* mouse administered with (right panel) or without (left panel) immune-depleting anti-CD8α antibodies. (**C**) Quantification of tumor infiltrating CD8^+^ and CD4^+^ T-cells from *MMTV-PyMT* mice treated for 28 days with 5-FU (40 mg/kg/4 days) with or without anti-CD8α antibodies (n=5-7, each dot represents an individual tumor and mouse), growth curves shown in Fig. 5J. Bar charts represent the mean and the dots show individual data points from individual tumors and mice. ** *P*<0.01, *** *P*<0.001.

## Notes

### Competing Interest Statement

The authors have declared no competing interest.

## References

1. A. Mantovani, F. Marchesi, A. Malesci, L. Laghi, P. Allavena, Tumour-associated macrophages as treatment targets in oncology. Nat Rev Clin Oncol 14, 399–416 (2017).

2. D. G. DeNardo et al., CD4(+) T cells regulate pulmonary metastasis of mammary carcinomas by enhancing protumor properties of macrophages. Cancer cell 16, 91–102 (2009).

3. T. Muliaditan et al., Macrophages are exploited from an innate wound healing response to facilitate cancer metastasis. Nature communications 9, 2951 (2018).

4. P. J. Murray et al., Macrophage activation and polarization: nomenclature and experimental guidelines. Immunity 41, 14–20 (2014).

5. R. Noy, J. W. Pollard, Tumor-Associated Macrophages: From Mechanisms to Therapy (vol 41, pg 49, 2014). Immunity 41, 866–866 (2014).

6. P. Paulus, E. R. Stanley, R. Schafer, D. Abraham, S. Aharinejad, Colony-stimulating factor-1 antibody reverses chemoresistance in human MCF-7 breast cancer xenografts. Cancer research 66, 4349–4356 (2006).

7. D. G. DeNardo et al., Leukocyte Complexity Predicts Breast Cancer Survival and Functionally Regulates Response to Chemotherapy. Cancer Discovery 1, 54–67 (2011).

8. B. Ruffell et al., Macrophage IL-10 blocks CD8+ T cell-dependent responses to chemotherapy by suppressing IL-12 expression in intratumoral dendritic cells. Cancer cell 26, 623–637 (2014).

9. R. A. Franklin et al., The cellular and molecular origin of tumor-associated macrophages. Science 344, 921–925 (2014).

10. O. R. Colegio et al., Functional polarization of tumour-associated macrophages by tumour-derived lactic acid. Nature 513, 559–563 (2014).

11. E. N. Arwert et al., A Unidirectional Transition from Migratory to Perivascular Macrophage Is Required for Tumor Cell Intravasation. Cell reports 23, 1239–1248 (2018).

12. Y. K. Huang et al., Macrophage spatial heterogeneity in gastric cancer defined by multiplex immunohistochemistry. Nature communications 10, 3928 (2019).

13. C. Carmona-Fontaine et al., Metabolic origins of spatial organization in the tumor microenvironment. Proceedings of the National Academy of Sciences of the United States of America 114, 2934–2939 (2017).

14. A. Lapenna, M. De Palma, C. E. Lewis, Perivascular macrophages in health and disease. Nature reviews. Immunology 18, 689–702 (2018).

15. C. E. Lewis, A. S. Harney, J. W. Pollard, The Multifaceted Role of Perivascular Macrophages in Tumors. Cancer Cell 30, 365 (2016).

16. J. W. Opzoomer et al., Macrophages orchestrate the expansion of a proangiogenic perivascular niche during cancer progression. Sci Adv 7, eabg9518 (2021).

17. J. B. Wyckoff et al., Direct visualization of macrophage-assisted tumor cell intravasation in mammary tumors. Cancer research 67, 2649–2656 (2007).

18. A. S. Harney et al., Real-Time Imaging Reveals Local, Transient Vascular Permeability, and Tumor Cell Intravasation Stimulated by TIE2hi Macrophage-Derived VEGFA. Cancer Discov 5, 932–943 (2015).

19. R. Hughes et al., Perivascular M2 Macrophages Stimulate Tumor Relapse after Chemotherapy. Cancer research 75, 3479–3491 (2015).

20. D. G. Jackson, R. Prevo, S. Clasper, S. Banerji, LYVE-1, the lymphatic system and tumor lymphangiogenesis. Trends Immunol 22, 317–321 (2001).

21. S. Ensan et al., Self-renewing resident arterial macrophages arise from embryonic CX3CR1(+) precursors and circulating monocytes immediately after birth. Nature immunology 17, 159–168 (2016).

22. C. H. Cho et al., Angiogenic role of LYVE-1-positive macrophages in adipose tissue. Circ Res 100, e47–57 (2007).

23. A. R. Pinto et al., An abundant tissue macrophage population in the adult murine heart with a distinct alternatively-activated macrophage profile. PLoS One 7, e36814 (2012).

24. H. Xu, M. Chen, D. M. Reid, J. V. Forrester, LYVE-1-positive macrophages are present in normal murine eyes. Invest Ophthalmol Vis Sci 48, 2162–2171 (2007).

25. S. Chakarov et al., Two distinct interstitial macrophage populations coexist across tissues in specific subtissular niches. Science 363, (2019).

26. C. Dollt et al., The shedded ectodomain of Lyve-1 expressed on M2-like tumor-associated macrophages inhibits melanoma cell proliferation. Oncotarget 8, 103682–103692 (2017).

27. M. De Palma et al., Tie2 identifies a hematopoietic lineage of proangiogenic monocytes required for tumor vessel formation and a mesenchymal population of pericyte progenitors. Cancer Cell 8, 211–226 (2005).

28. C. T. Guy, R. D. Cardiff, W. J. Muller, Induction of mammary tumors by expression of polyomavirus middle T oncogene: a transgenic mouse model for metastatic disease. Mol Cell Biol 12, 954–961 (1992).

29. S. Z. Wu et al., A single-cell and spatially resolved atlas of human breast cancers. Nat Genet 53, 1334–1347 (2021).

30. R. Gozzelino, V. Jeney, M. P. Soares, Mechanisms of cell protection by heme oxygenase-1. Annu Rev Pharmacol Toxicol 50, 323–354 (2010).

31. K. N. Luu Hoang, J. E. Anstee, J. N. Arnold, The Diverse Roles of Heme Oxygenase-1 in Tumor Progression. Front Immunol 12, 658315 (2021).

32. J. N. Arnold, L. Magiera, M. Kraman, D. T. Fearon, Tumoral immune suppression by macrophages expressing fibroblast activation protein-alpha and heme oxygenase-1. Cancer immunology research 2, 121–126 (2014).

33. Q. Tan et al., Src/STAT3-dependent heme oxygenase-1 induction mediates chemoresistance of breast cancer cells to doxorubicin by promoting autophagy. Cancer Sci 106, 1023–1032 (2015).

34. P. Nuhn et al., Heme oxygenase-1 and its metabolites affect pancreatic tumor growth in vivo. Mol Cancer 8, 37 (2009).

35. D. Nowis et al., Zinc protoporphyrin IX, a heme oxygenase-1 inhibitor, demonstrates potent antitumor effects but is unable to potentiate antitumor effects of chemotherapeutics in mice. BMC Cancer 8, 197 (2008).

36. S. Di Biase et al., Fasting-Mimicking Diet Reduces HO-1 to Promote T Cell-Mediated Tumor Cytotoxicity. Cancer Cell 30, 136–146 (2016).

37. P. O. Berberat et al., Inhibition of heme oxygenase-1 increases responsiveness of pancreatic cancer cells to anticancer treatment. Clinical Cancer Research 11, 3790–3798 (2005).

38. P. Kosti et al., Hypoxia-sensing CAR T cells provide safety and efficacy in treating solid tumors. Cell Rep Med 2, 100227 (2021).

39. J. H. Kim et al., High cleavage efficiency of a 2A peptide derived from porcine teschovirus-1 in human cell lines, zebrafish and mice. PLoS One 6, e18556 (2011).

40. S. Spranger, D. Dai, B. Horton, T. F. Gajewski, Tumor-Residing Batf3 Dendritic Cells Are Required for Effector T Cell Trafficking and Adoptive T Cell Therapy. Cancer Cell 31, 711–723 e714 (2017).

41. M. Haldar et al., Heme-mediated SPI-C induction promotes monocyte differentiation into iron-recycling macrophages. Cell 156, 1223–1234 (2014).

42. M. Tomura et al., Monitoring cellular movement in vivo with photoconvertible fluorescence protein “Kaede” transgenic mice. Proceedings of the National Academy of Sciences of the United States of America 105, 10871–10876 (2008).

43. A. Kramer, J. Green, J. Pollard, Jr., S. Tugendreich, Causal analysis approaches in Ingenuity Pathway Analysis. Bioinformatics 30, 523–530 (2014).

44. H. Moura Silva et al., c-MAF-dependent perivascular macrophages regulate diet-induced metabolic syndrome. Sci Immunol 6, eabg7506 (2021).

45. D. E. Johnson, R. A. O’Keefe, J. R. Grandis, Targeting the IL-6/JAK/STAT3 signalling axis in cancer. Nat Rev Clin Oncol 15, 234–248 (2018).

46. A. Sahoo et al., Batf is important for IL-4 expression in T follicular helper cells. Nature communications 6, 7997 (2015).

47. C. D. Mills, K. Kincaid, J. M. Alt, M. J. Heilman, A. M. Hill, M-1/M-2 macrophages and the Th1/Th2 paradigm. Journal of immunology 164, 6166–6173 (2000).

48. G. Fatkenheuer et al., Efficacy of short-term monotherapy with maraviroc, a new CCR5 antagonist, in patients infected with HIV-1. Nat Med 11, 1170–1172 (2005).

49. D. Vestweber, How leukocytes cross the vascular endothelium. Nature reviews. Immunology 15, 692–704 (2015).

50. R. J. Wong et al., In vitro inhibition of heme oxygenase isoenzymes by metalloporphyrins. Journal of perinatology: official journal of the California Perinatal Association 31 Suppl 1, S35–41 (2011).

51. P. C. Tumeh et al., PD-1 blockade induces responses by inhibiting adaptive immune resistance. Nature 515, 568–571 (2014).

52. O. Hamid et al., A prospective phase II trial exploring the association between tumor microenvironment biomarkers and clinical activity of ipilimumab in advanced melanoma. J Transl Med 9, 204 (2011).

53. M. Hong et al., Chemotherapy induces intratumoral expression of chemokines in cutaneous melanoma, favoring T-cell infiltration and tumor control. Cancer research 71, 6997–7009 (2011).

54. S. Ladoire et al., In situ immune response after neoadjuvant chemotherapy for breast cancer predicts survival. J Pathol 224, 389–400 (2011).

55. T. Muliaditan et al., Repurposing Tin Mesoporphyrin as an Immune Checkpoint Inhibitor Shows Therapeutic Efficacy in Preclinical Models of Cancer. Clinical cancer research: an official journal of the American Association for Cancer Research 24, 1617–1628 (2018).

56. D. B. Longley, D. P. Harkin, P. G. Johnston, 5-fluorouracil: mechanisms of action and clinical strategies. Nature reviews. Cancer 3, 330–338 (2003).

57. S. Ugel et al., Immune tolerance to tumor antigens occurs in a specialized environment of the spleen. Cell reports 2, 628–639 (2012).

58. Z. Cao et al., Antitumor and immunomodulatory effects of low-dose 5-FU on hepatoma 22 tumor-bearing mice. Oncol Lett 7, 1260–1264 (2014).

59. A. F. Welford et al., TIE2-expressing macrophages limit the therapeutic efficacy of the vascular-disrupting agent combretastatin A4 phosphate in mice. J Clin Invest 121, 1969–1973 (2011).

60. A. S. Harney et al., The Selective Tie2 Inhibitor Rebastinib Blocks Recruitment and Function of Tie2(Hi) Macrophages in Breast Cancer and Pancreatic Neuroendocrine Tumors. Mol Cancer Ther 16, 2486–2501 (2017).

61. R. Mazzieri et al., Targeting the ANG2/TIE2 axis inhibits tumor growth and metastasis by impairing angiogenesis and disabling rebounds of proangiogenic myeloid cells. Cancer Cell 19, 512–526 (2011).

62. S. W. Ryter, A. M. Choi, Heme oxygenase-1/carbon monoxide: from metabolism to molecular therapy. American journal of respiratory cell and molecular biology 41, 251–260 (2009).

63. N. Casares et al., Caspase-dependent immunogenicity of doxorubicin-induced tumor cell death. The Journal of experimental medicine 202, 1691–1701 (2005).

64. S. M. Geary, C. D. Lemke, D. M. Lubaroff, A. K. Salem, The combination of a low-dose chemotherapeutic agent, 5-fluorouracil, and an adenoviral tumor vaccine has a synergistic benefit on survival in a tumor model system. PLoS One 8, e67904 (2013).

65. L. Bracci, G. Schiavoni, A. Sistigu, F. Belardelli, Immune-based mechanisms of cytotoxic chemotherapy: implications for the design of novel and rationale-based combined treatments against cancer. Cell Death Differ 21, 15–25 (2014).

66. C. Pfirschke et al., Immunogenic Chemotherapy Sensitizes Tumors to Checkpoint Blockade Therapy. Immunity 44, 343–354 (2016).

67. J. W. Opzoomer, D. Sosnowska, J. E. Anstee, J. F. Spicer, J. N. Arnold, Cytotoxic Chemotherapy as an Immune Stimulus: A Molecular Perspective on Turning Up the Immunological Heat on Cancer. Front Immunol 10, 1654 (2019).

68. A. Sistigu et al., Cancer cell-autonomous contribution of type I interferon signaling to the efficacy of chemotherapy. Nat Med 20, 1301–1309 (2014).

69. E. H. Bent et al., Microenvironmental IL-6 inhibits anti-cancer immune responses generated by cytotoxic chemotherapy. Nature communications 12, 6218 (2021).

70. L. E. Otterbein et al., Carbon monoxide has anti-inflammatory effects involving the mitogen-activated protein kinase pathway. Nature medicine 6, 422–428 (2000).

71. X. Zhang, P. Shan, J. Alam, X. Y. Fu, P. J. Lee, Carbon monoxide differentially modulates STAT1 and STAT3 and inhibits apoptosis via a phosphatidylinositol 3-kinase/Akt and p38 kinase-dependent STAT3 pathway during anoxia-reoxygenation injury. The Journal of biological chemistry 280, 8714–8721 (2005).

72. G. Cepinskas, K. Katada, A. Bihari, R. F. Potter, Carbon monoxide liberated from carbon monoxide-releasing molecule CORM-2 attenuates inflammation in the liver of septic mice. Am J Physiol Gastrointest Liver Physiol 294, G184–191 (2008).

73. J. Megias, J. Busserolles, M. J. Alcaraz, The carbon monoxide-releasing molecule CORM-2 inhibits the inflammatory response induced by cytokines in Caco-2 cells. Br J Pharmacol 150, 977–986 (2007).

74. J. J. Muldoon, Y. Chuang, N. Bagheri, J. N. Leonard, Macrophages employ quorum licensing to regulate collective activation. Nature communications 11, 878 (2020).

75. S. Goswami et al., Identification of invasion specific splice variants of the cytoskeletal protein Mena present in mammary tumor cells during invasion in vivo. Clin Exp Metastasis 26, 153–159 (2009).

76. M. Roh-Johnson et al., Macrophage contact induces RhoA GTPase signaling to trigger tumor cell intravasation. Oncogene 33, 4203–4212 (2014).

77. B. D. Robinson et al., Tumor microenvironment of metastasis in human breast carcinoma: a potential prognostic marker linked to hematogenous dissemination. Clinical cancer research: an official journal of the American Association for Cancer Research 15, 2433–2441 (2009).

78. T. E. Rohan et al., Tumor microenvironment of metastasis and risk of distant metastasis of breast cancer. J Natl Cancer Inst 106, (2014).

79. J. A. Sparano et al., A metastasis biomarker (MetaSite Breast Score) is associated with distant recurrence in hormone receptor-positive, HER2-negative early-stage breast cancer. NPJ Breast Cancer 3, 42 (2017).

80. A. Abtin et al., Perivascular macrophages mediate neutrophil recruitment during bacterial skin infection. Nature immunology 15, 45–53 (2014).

81. N. Halama et al., Tumoral Immune Cell Exploitation in Colorectal Cancer Metastases Can Be Targeted Effectively by Anti-CCR5 Therapy in Cancer Patients. Cancer Cell 29, 587–601 (2016).

## Supplementary References

82. S. Tzima, P. Victoratos, K. Kranidioti, M. Alexiou, G. Kollias, Myeloid heme oxygenase-1 regulates innate immunity and autoimmunity by modulating IFN-beta production. The Journal of experimental medicine 206, 1167–1179 (2009).

83. W. C. Skarnes et al., A conditional knockout resource for the genome-wide study of mouse gene function. Nature 474, 337–342 (2011).

84. M. Kraman et al., Suppression of antitumor immunity by stromal cells expressing fibroblast activation protein-alpha. Science 330, 827–830 (2010).

85. E. E. Dutton et al., Peripheral lymph nodes contain migratory and resident innate lymphoid cell populations. Sci Immunol 4, (2019).

86. B. Z. Qian et al., CCL2 recruits inflammatory monocytes to facilitate breast-tumour metastasis. Nature 475, 222–225 (2011).

87. T. Stuart et al., Comprehensive Integration of Single-Cell Data. Cell 177, 1888–1902 e1821 (2019).

88. H. A. Pliner, J. Shendure, C. Trapnell, Supervised classification enables rapid annotation of cell atlases. Nat Methods 16, 983–986 (2019).

89. A. Sharma et al., Onco-fetal Reprogramming of Endothelial Cells Drives Immunosuppressive Macrophages in Hepatocellular Carcinoma. Cell 183, 377–394 e321 (2020).

90. B. M. Bolstad, R. A. Irizarry, M. Astrand, T. P. Speed, A comparison of normalization methods for high density oligonucleotide array data based on variance and bias. Bioinformatics 19, 185–193 (2003).

91. C. M. Elso et al., Leishmaniasis host response loci (lmr1-3) modify disease severity through a Th1/Th2-independent pathway. Genes Immun 5, 93–100 (2004).

